# Alu-mediated weak CEBPA binding and slow B cell transdifferentiation in human

**DOI:** 10.1101/2021.10.28.466072

**Authors:** Ramil Nurtdinov, María Sanz, Amaya Abad, Alexandre Esteban, Sebastian Ullrich, Carme Arnan, Rory Johnson, Sílvia Pérez-Lluch, Roderic Guigó

## Abstract

Many developmental and differentiation processes take substantially longer in human than in mouse. To investigate the molecular mechanisms underlying this phenomenon, here we have specifically focused on the transdifferentiation from B cells to macrophages. The process is triggered by exactly the same molecular mechanism -- the induction by the transcription factor (TF) CEBPA -- but takes three days in mouse and seven in human (*1, 2*). In mouse, the speed of this process is known to be associated with *Myc* expression (*3*). We found that in this species, CEBPA binds strongly to the *Myc* promoter, efficiently down-regulating *Myc*. In human, in contrast, CEBPA does not bind this promoter, and *MYC* is indirectly and more slowly down-regulated. Attenuation of CEBPA binding is not specific to the *MYC* promoter, but a general trait of the human genome across multiple biological conditions. We traced back weak CEBPA binding to the primate-specific Alu repeat expansion. Many Alu repeats carry strong CEBPA binding motifs, which sequester CEBPA, and attenuate CEBPA binding genome-wide. We observed similar CEBPA and *MYC* dynamics in natural processes regulated by CEBPA, suggesting that CEBPA attenuation could underlie the longer duration in human processes controlled by this factor. Our work highlights the highly complex mode in which biological information is encoded in genome sequences, evolutionarily connecting, in an unexpected way, lineage-specific transposable element expansions to species-specific changes in developmental tempos.

## Main

To understand the differences in duration between human and mouse B cell transdifferentiation after CEBPA induction, we performed RNA-Seq and CEBPA ChIP-Seq at 12 time-points in human BLaER1 cells, and RNA-Seq at 10 time-points in mouse C10 cells (*1, 2*) and reanalyzed published CEBPA ChIP-Seq in these cells (*4*) (Extended Data Fig. 1a, Supplementary Table 1, Methods). For genes up- or down-regulated during transdifferentiation -- the most likely to be involved in the transition between cell types (Extended Data Fig. 1b, Extended Data Fig. 2) -- we calculated the speed of regulation as the time at which expression changes two-fold from the initial expression (Extended Data Fig. 1c, Methods). Both up- and, in particular, down-regulation are delayed in human compared to mouse (Fig. 1a-b, Extended Data Fig. 1d).

**Fig. 1.**
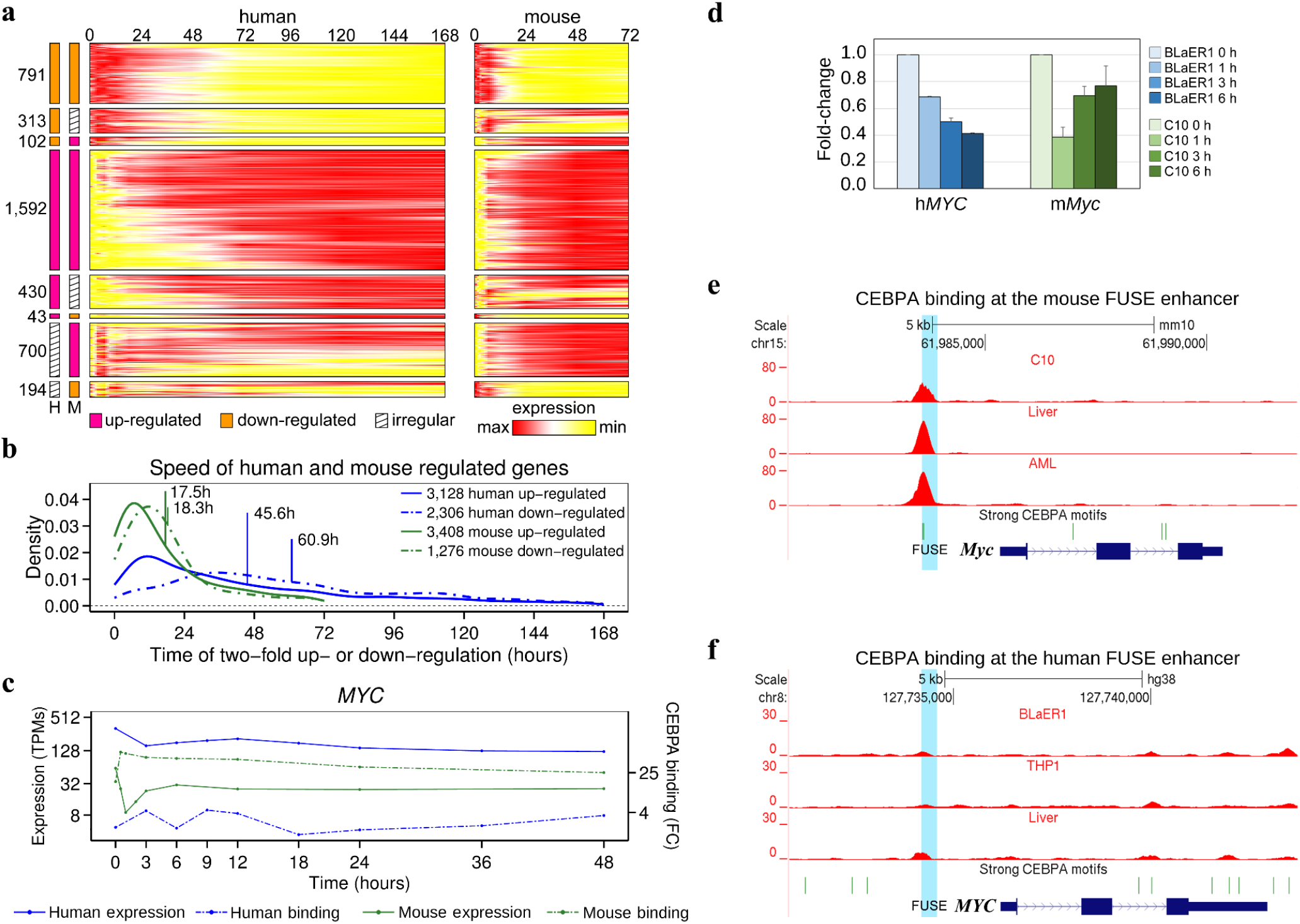
Transcriptional dynamics during transdifferentiation. **a,** Expression profiles during transdifferentiation of orthologous human and mouse genes. Only up- and down-regulated genes in at least one species are shown. Expression values are normalized between 0 (minimal expression) and 1 (maximum expression) within each gene independently. The number of genes in each category is shown at the left. **b,** Distribution of the speed of up- and down-regulation in genes regulated during transdifferentiation (see text). Means for each distribution are given. **c,** Expression profiles for human and mouse *MYC* expression and CEBPA binding at the FUSE enhancer. Maximum *MYC* downregulation is at 1h post-induction (9 TPMs) in mouse, and at 120h (43 TPMs) in human. **d,** Relative *MYC* expression in human and mouse 0 h, 1 h, 3 h and 6h after transdifferentiation induction measured by quantitative PCR. Four biological replicates were performed for each cell type. Error bars represent Standard Error of the Mean (SEM). **e,** Fold-change signal tracks for CEBPA binding at the *Myc* locus in mouse C10 cells three hours after induction, liver and AML cells. **f,** Fold-change signal tracks for CEBPA binding at the *MYC* locus in human BLaER1 cells three hours after induction, liver and THP1 cells.

Among down-regulated genes we found *MYC* (Fig. 1c). It has been recently shown that *MYC* influences the speed of the transdifferentiation in mouse: cells with low *Myc* activity transdifferentiate into macrophages very efficiently, while cells with high *Myc* activity transdifferentiate inefficiently (*3*). During transdifferentiation, *Myc* responds to CEBPA induction very rapidly in mouse C10 cells (minimum expression 1 hour after induction) whereas in human BLaER1 cells, where *MYC* expression levels are consistently higher, downregulation is substantially delayed (minimum expression at 120 hours with a local minimum between 3 and 6 hours, Fig. 1c-d, Extended Data Fig. 3a). A large fraction of genes regulated during transdifferentiation are candidate MYC targets both in human and mouse, a trend that is even stronger for down-regulated genes (Extended Data Fig. 3d, data from (*5–7*), Methods). All this together suggests that constitutively high expression of *MYC* and inefficient inactivation by CEBPA contributes importantly to the delay, in human cells, of the cascade of molecular events responsible for transdifferentiation.

We, thus, analyzed CEBPA binding at the far upstream sequence element (FUSE) enhancer known to regulate *MYC* expression (*8, 9*). In C10 cells, induction of transdifferentiation results in strong CEBPA binding, (Fig. 1e, Extended Data Fig. 3a,c). In contrast, in human BLaER1 binding is very weak (Fig. 1f, Extended Data Fig. 3a,b). Overall there is a strong negative correlation between CEBPA binding and *Myc* expression (cc=-0.71, p-val=0.077) in mouse C10 cells during transdifferentiation, while there is essentially no correlation (cc=0.31, p-val=0.322) in BLaER1 cells. The dynamics of the relationship between CEBPA binding and *MYC* expression is conserved also along mouse liver regeneration, a natural process regulated by CEBPA (*10–12*). Liver regenerating after partial hepatectomy shows reduced CEBPA binding and increased *Myc* expression, when compared with normal liver (Extended Data Fig. 3e).

Underlying the CEBPA peaks at FUSE, we found two CEBPA binding sites, which are stronger in the mouse genome because of mutations at the +/- 4 positions, which have apparently arosen in the muroidea lineage (Extended Data Fig. 4c). These sites are the most critical determinants of CEBPA recognition specificity and affinity (*13*), and we took advantage of the mutations at these sites to demonstrate that CEBPA binding at FUSE efficiently down-regulates *Myc* in mouse, but that in human, in contrast, *MYC* down-regulation is largely independent of direct CEBPA binding. We designed reporter assays in which the expression of the reporter gene *mCherry* is under the control of 2-kb *MYC* promoters either from mouse or human, including the FUSE and the CEBPA binding motifs. In parallel, we generated reporter constructs substituting the CEBPA binding sites in the mouse *Myc* promoter for those in human, and vice versa (Fig. 2a).

**Fig. 2.**
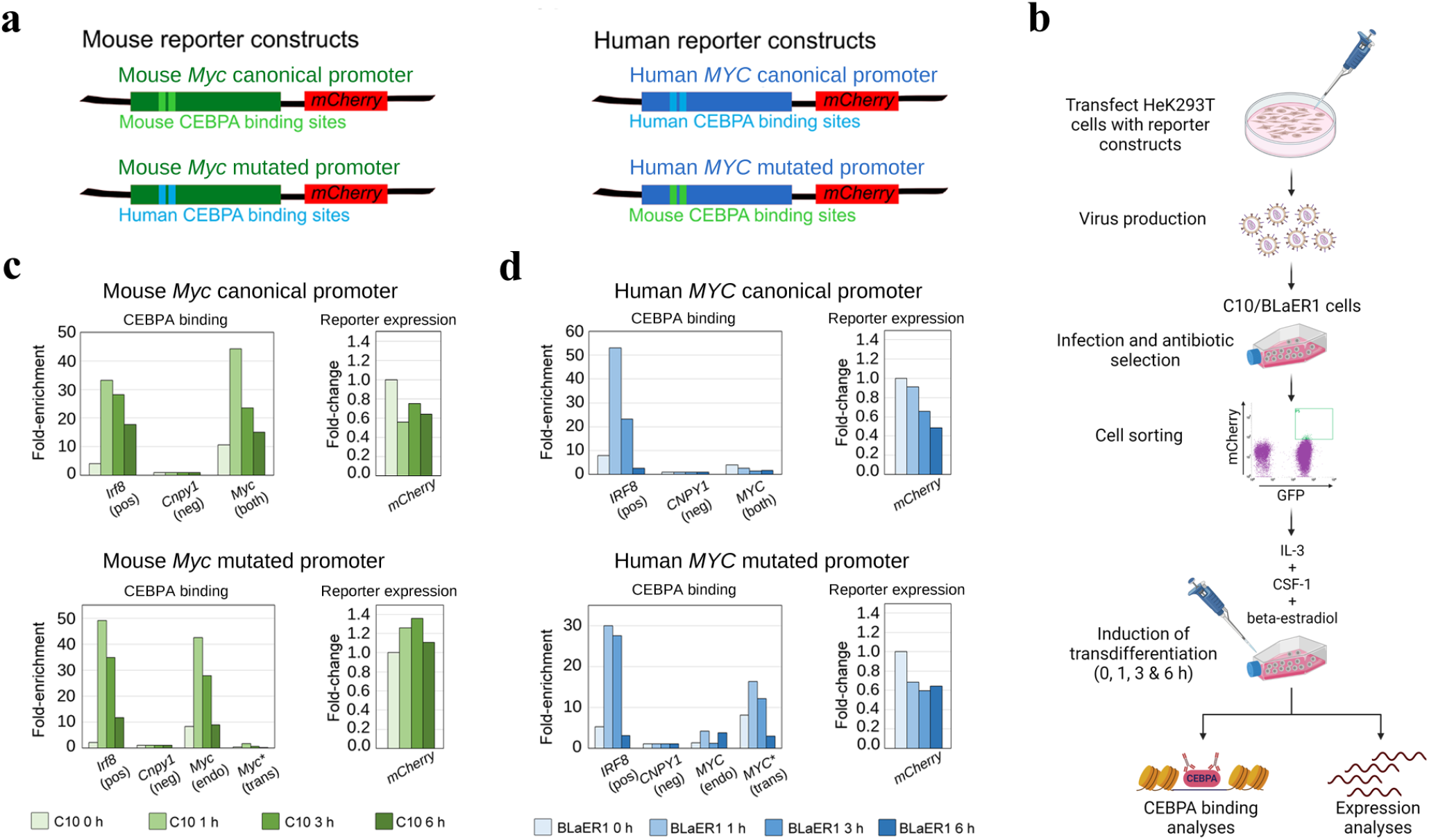
Disruption of CEBPA binding impairs down-regulation in mouse C10 cells. **a,** Reporter constructs expressing the *mCherry* reporter gene under the control of the 2-Kb promoter region of the mouse *Myc* (left, green boxes) and the human *MYC* (right, blue boxes) gene, including the FUSE enhancer, which includes the CEBPA binding sites (highlighted as lighter boxes). In both cases, constructs containing the mutated CEBPA binding sites were also generated. Two point mutations were introduced in the mouse reporter construct to revert mouse CEBPA binding sites to human sequences, and vice versa. **b,** Experimental workflow of the reporter assays. HeK293T virus packaging cells were transfected with the four constructs in **a** and C10 and BLaER1 cells were infected with viruses containing the different constructs. After antibiotic selection, cells positive for GFP (co-expressed with CEBPA) and mCherry (included in the reporter construct) markers were sorted by FACS. Transdifferentiation was induced and cells were collected at time-points 0, 1, 3 and 6 h after induction. From each time-point, chromatin and RNA were extracted to perform CEBPA binding and expression analyses, respectively. **c,** Reporter experiments in mouse C10 cells. Upper panels, CEBPA binding (left) and expression of the reporter gene *mCherry* (right) in cells infected with the construct carrying the canonical mouse *Myc* promoter. Lower panels, CEBPA binding (left) and expression of the reporter gene *mCherry* (right) in cells infected with the construct carrying the mutated mouse *Myc* promoter. **d,** Reporter experiments in human BLaER1 cells. Upper panels, CEBPA binding (left) and expression of the reporter gene *mCherry* (right) in cells infected with the construct carrying the canonical human *MYC* promoter. Lower panels, CEBPA binding (left) and expression of the reporter gene *mCherry* (right) in cells infected with the construct carrying the mutated human *MYC* promoter. In **c** and **d**, *IRF8* and *CNPY1* are used as positive (pos) and negative (neg) controls, respectively of CEBPA binding. For *MYC* promoters, primers amplifying the endogenous copy (endo), the transgenic copy (trans) or both copies in parallel (both) were used. At least, two biological replicates were performed per experiment.

We monitored CEBPA binding, and the expression of the reporter *mCherry* before and after induction of transdifferentiation in BLaER1 and C10 cells infected with these reporters (Fig. 2b). Mouse C10 cells infected with the canonical mouse promoter show strong CEBPA binding, and abrupt down-regulation of the *mCherry* reporter at 1h followed by mild recovery afterwards (Fig. 2c, upper panels), as observed for endogenous *Myc* (Fig. 1d). Binding is impaired, in contrast, in C10 cells infected with the promoter in which the mouse CEBPA binding motifs have been converted to the human sequence, and there is no down-regulation of *mCherry* after induction (Fig. 2c, lower panels).

In BLaER1 cells infected with the canonical human promoter, there is no recruitment of CEBPA and, as with endogenous *MYC* (Fig. 1d), we observed down-regulation of *mCherry* after induction (Fig. 2d, upper panels). Recruitment of CEBPA is induced in BLaER1 cells infected with the promoter carrying the mouse CEBPA binding motif. However, CEBPA binding has, in this case, little impact on *mCherry* expression (Fig. 2d, lower panels).

Weaker CEBPA binding in human than mouse is not specific to the *MYC* enhancer during transdifferentiation, but a general trait of the human genome across multiple biological conditions: CEBPA binds globally more strongly to the mouse than to the human genome. This can be observed when comparing matched time-points during transdifferentiation (Fig. 3a, Extended Data Fig. 5a-b, Supplementary Methods), when analyzing comparable cell types in human and mouse: liver (*15*), and human THP1 (*16*) and mouse AML (*17*) cells (Fig. 3a) and, more generally, when analyzing dozens of human and mouse CEBPA ChIP-Seq data across multiple cell types and tissues uniformly processed at the ChIP-Atlas database (*18*) (Fig. 3b).

**Fig. 3.**
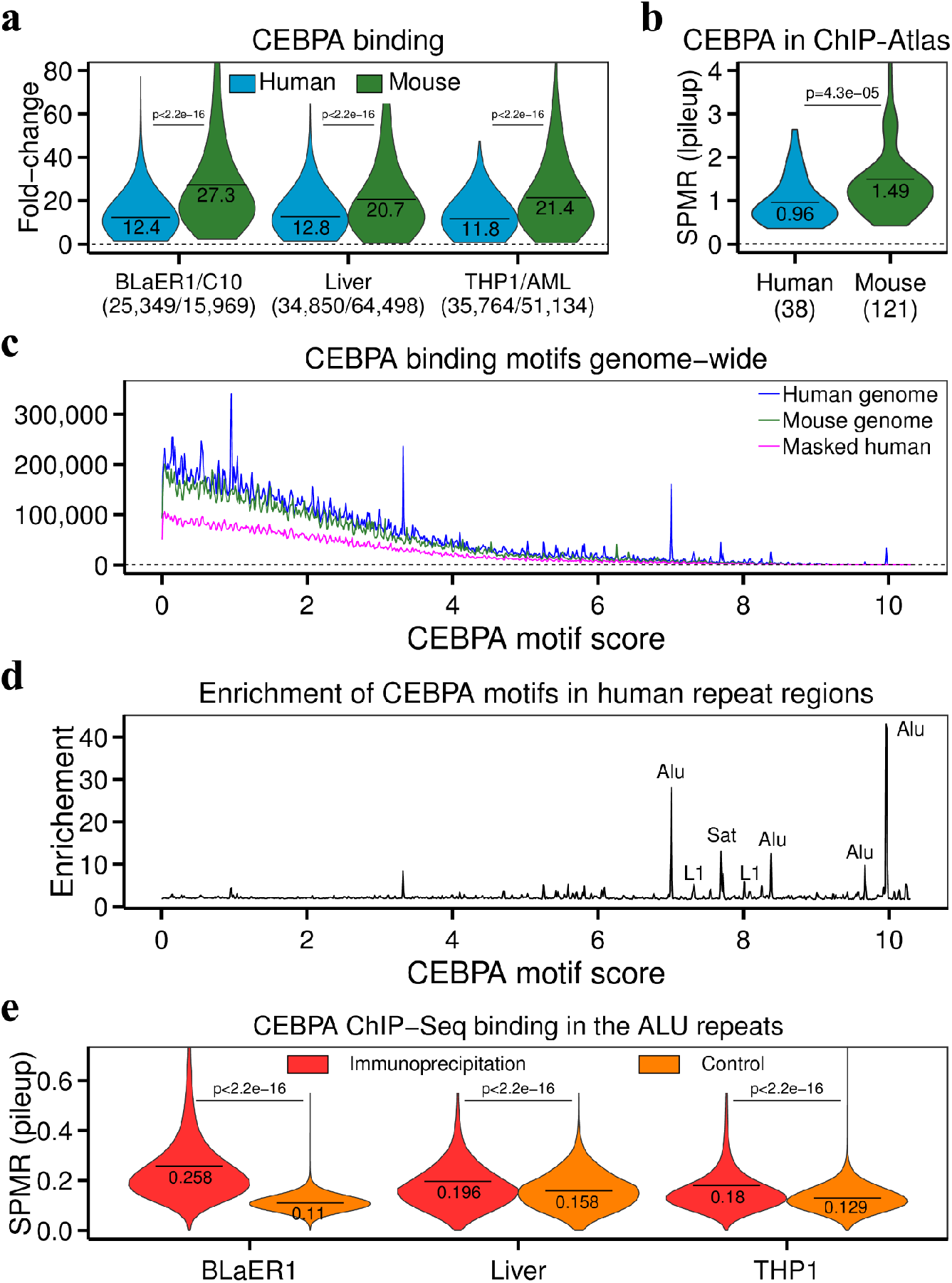
CEBPA binding sites in Alu repeats. **a,** Distribution of fold-change of CEBPA peaks genome-wide in human and mouse. For BLaER1/C10 the values correspond to those at 3h after induction. The values within brackets correspond to the number of peaks identified within a particular sample. **b,** Distribution of average CEBPA peak strength (SPMR) across CEBPA ChIP-Seq samples in ChIP-Atlas. The values within brackets correspond to the number of samples in ChIP-Atlas. **c,** Distribution of the strength of CEBPA binding sites within human and mouse genomes. The score of the binding sites has been computed using the human CEBPA PWM from Hocomoco(*14*). See Extended Data Fig. 5c for the distribution using the mouse PWM. **d,** Enrichment of CEBPA binding sites in human repeated regions. For each PWM score, the enrichment is the number of matches of the CEBPA motif with this score in the unmasked genome over the number of matches in the masked genome. We highlighted the type of repeat for scores greater than 7 and enrichments greater than 6. **e,** Distribution of pileup signal (SPMR) in Alu regions using CEBPA ChIP-Seq and input data for BLaER1 cells, liver and THP1. In **a**, **b** and **e**, horizontal lines represent the mean value.

To understand the sequence determinants underlying the weaker CEBPA binding in the human genome, we searched for CEBPA binding motifs in the human and mouse genomes using a standard CEBPA Position Weight Matrix. We found many more strong CEBPA binding motifs in the human than in the mouse genome (Fig. 3c). This originates, almost exclusively, from the overrepresentation of a few specific instances of the CEBPA consensus motif (Fig. 3d, Extended Data Fig. 5c) which are located within about 400,000 repetitive regions, mostly Alu (Supplementary Table 1). There is no overrepresentation of CEBPA binding motifs in mouse repetitive regions (Extended Data Fig. 5c).

We hypothesized that the expansion of CEBPA binding sites in the human genome leads to an increase in the competition for CEBPA binding. Alu sequences would sequester a large fraction of CEBPA molecules, resulting in the overall weaker CEBPA binding observed in the human compared to the mouse genome. Indeed, we found that Alu repeats containing strong binding sites attract a much larger number of CEBPA ChIP-Seq reads than expected (Fig. 3e, Supplementary Table 3, Methods). As a consequence, equally strong sequence binding motifs attract CEBPA more strongly in mouse than in human (Extended Data Fig. 5d). Conservative simulations suggest that Alu competition for CEBPA binding could explain at least 30% of the difference in CEBPA binding observed between human and mouse during transdifferentiation (Supplementary Table 3, Supplementary Methods).

Overall attenuation of CEBPA binding is likely to have an impact on longer transdifferentiation, beyond that mediated by MYC. Indeed, we found a weak, but significant, negative correlation between strength of the binding and speed of regulation of potential CEBPA targets regulated during transdifferentiation (Methods), both in human (cc=-0.11, p-val=0.0003), and mouse (cc=-0.14, p-val=8.0e-07). More importantly, we specifically observed weak CEBPA binding in other key transdifferentiation factors. For instance *SPI1*, the master regulator promoting the expression of macrophage specific genes (*19*) and known to be activated by CEBPA (*20*), shows delayed up-regulation and some weaker CEBPA binding in human compared mouse, even though the underlying regulatory sequence is identical in the two species (Extended Data Fig. 6). On the other hand, *PAX5*, the repression of which is necessary for transdifferentiation, as it maintains the expression of B cell specific genes (*21*), shows a pattern similar to that of *MYC*. Down-regulation of *PAX5* is strongly delayed in human compared to mouse, and while there is strong CEBPA binding at the *PAX5* regulatory regions in mouse C10 cells, there is no binding in human BLaER1 cells (Extended Data Fig. 7). We found, however, that *ID1* and *ID2*, known repressors of *PAX5* (*22, 23*), are up-regulated in both human and mouse, and show strong CEBPA binding in the two species (Extended Data Fig. 8, Fig. 9), suggesting a mechanism of indirectly regulation of *PAX5* by CEBPA.

A similar indirect mechanism, mediated by CEBPA-induced activation of repressors, could explain the observed down-regulation of *MYC* in human BLaER1 cells. For instance, one notable exception in which there is strong CEBPA binding in human, but very weak in mouse is *PRDM1*, a known repressor of *MYC* and *PAX5* (*24–26*). Consistently, in mouse C10 cells, *PRDM1* remains very lowly expressed, but it is strongly up-regulated in BLaER1 cells upon induction (Extended Data Fig. 10).

## Discussion

In contrast to previous reports, in which the longer duration of physiological processes in human than in mouse has been attributed to generic slower biochemical reactions (*27*) and/or longer protein stability (*28*), here we propose that slower B cell transdifferentiation can be, at least partially, attributed to a specific cause: the attenuation of the binding of a TF to the human genome. The little comparative data currently available suggest that general unresponsiveness to CEBPA could generally lead to longer duration in human of the physiological processes controlled by this factor. Indeed, it has been recently shown that during liver embryogenesis, where CEBPA plays a role (*10*), hepatocytes differentiate from hepatoblasts in about three weeks in humans, but only in five days in mice (*29*). It is unclear whether duration of individual physiological processes relates to organismic life-span. It is remarkable, however, that reduced expression of CEBPB, another key hepatocyte transcription factor with which CEBPA shares substantial sequence similarity, has been recently shown to increase life-span in mouse (*30*).

Repeat-associate expansion of TF binding motifs along genomes have been previously reported, as in the SINE-associated expansion of CTCF binding sites in the human genome (*31, 32*), or the transposable element-mediated duplication of REST binding sites (*33*) in mammals. Here we show that this expansion has an important functional impact by globally attenuating the response to a particular TF. This attenuation does not appear to be the result of direct selection, but the unpredictable by-product of evolution at genome scale. The issue of the fraction of the genome that carries biological functionality has stirred heated debates (*34–36*). It is often assumed that proof of selection is the ultimate demonstration of biological functionality. Our results show, however, that, given the highly interconnected and epistatic nature of genomic information, evolutionary processes with profound biological implications, may not leave the direct imprint of selection.

## Methods

### Induction of transdifferentiation and flow cytometry

Human BLaER1 and mouse C10 cell lines were maintained and transdifferentiation was induced as previously reported (*1, 2*). Fluorescent staining of cell surface markers was done with antibodies against Mac-1 (APC) and CD19 (APC-Cy7) (BD Pharmigen). Samples were monitored on the LSRII flow cytometer (BD Biosciences) and data were analyzed with FlowJo software (Tree Star).

### RNA extraction, retrotranscription, library preparation and sequencing

For RNA-Seq, RNA samples were collected at 0, 0.5, 1, 2, 3, 6, 12, 24, 48 and 72 hours after transdifferentiation induction in C10 mouse cells, and at 0, 3, 6, 9, 12, 18, 24, 36, 48, 72, 120 and 168 hours after transdifferentiation induction in BLaER1 human cells, to allow for a maximum resolution of the process, following the original reports (*1, 2*). Briefly, cells were lysed with Qiazol following the manufacturer’s instructions (Qiagen). RNA was isolated and purified by using RNeasy mini kit columns, also following the manufacturer’s instructions (Qiagen). Stranded poly A+ libraries were prepared with 1 μg of total RNA using TruSeq mRNA Sample Prep Kit (Illumina) according to the manufacturer’s protocol. Libraries were analyzed using Agilent DNA 1000 chips to determine the quantity and size distribution and sequenced on Illumina’s HiSeq2000.

For *MYC* and reporter expression analyses, RNA from 500,000 BLaER1 and 750,000 C10 cells at 0, 1, 3, and 6 hours after transdifferentiation induction was extracted with Zymo RNA Miniprep kit, following the manufacturer’s instructions. RNA was retrotranscribed with RevertAid retrotranscriptase (Thermo) and gene expression was assessed by quantitative PCR in a Roche LightCycler480. Expression of *MYC* and the reporter gene was assessed by relative quantification to *GAPDH* expression and to time point 0 h after induction (delta-delta-Ct method), and normalizing by the amplification efficiency of each primer set. Primers used for qPCR are listed in Supplementary Methods. At least, two biological replicates were performed for each experiment.

### Chromatin immunoprecipitation, library preparation and sequencing

For ChIP-Seq experiments, cells were crosslinked with 1% formaldehyde for 10 minutes at room temperature and sheared with a Covaris sonicator (ChIP-Seq) or with a Diagenode Bioruptor or a Qsonica sonicator (individual ChIPs). For ChIP-Seq, 5 μg of chromatin and 5 μg of antibody against rat CEBPA (sc-61 X, Santa Cruz) were incubated in RIPA buffer (140 mM NaCl, 10 mM Tris HCl pH 8.0, 1 mM EDTA, 1 % Triton X-100, 0.1 % SDS, 0.1 % Na deoxycholate, protease inhibitors). ChIP-Seq libraries were prepared with 1 ng of purified ChIP using NEBNext DNA Library kit for Illumina following the manufacturer’s protocol. Libraries were sequenced on an Illumina’s HiSeq2000 machine. For individual ChIPs, 1 μg of chromatin was incubated with 1 μg of antibody in the same conditions as above. Individual ChIPs were analyzed by quantitative PCR in a Roche LightCycler480. Primers used for qPCR are listed in Supplementary Methods. At least, two biological replicates were performed both for ChIP-Seq and for individual ChIPs.

### Processing of RNASeq

We mapped our pair-end reads to human (hg38) and mouse (mm10) genomes as well as human (Gencode v30) and mouse (Gencode vM22) transcriptomes (*37*) with STAR aligner (*38*). Approximately 90% human and 82% mouse reads pairs were mapped uniquely to corresponding genomes, 5% human and 11% mouse reads were mapped in multiple locations. 81% human and 80% mouse reads were mapped to corresponding transcriptomes and were used to quantify gene and transcript expression, as TPM (Transcripts Per Million) with RSEM (*39*) (Supplementary Table 1).

### Principal Component analysis and gene expression heatmaps of human and mouse samples

We used Ensembl BioMart (*40*) to obtain a list of 16,040 orthologous human-mouse protein-coding genes. We set a threshold of 0.1 TPMs in at least two time-points in human and mouse to consider a gene expressed in both species. Thus we obtained a set of 9,188 expressed orthologous human-mouse protein-coding genes. We merged 12 human and 10 mouse expression values for each orthologous gene and applied z-score normalization of the resulting expression vectors by subtracting the mean across all 22 values and dividing by standard deviation. The resulting matrix was subjected to PCA analysis with prcomp R function (*41*). We plot the values of the two first principal components at Extended Data Fig. 1.

To create the expression heatmaps for orthologous protein-coding genes we normalized the expression profile of each gene independently. We subtracted from the expression of each gene at each time-point the minimal expression of the gene across all time-points and divided by the difference between maximal and minimal expression, thus fitting the resulting profile into 0:1 interval. Next we imputed expression at each hour using linear interpolation and obtained 73 normalized expression values for mouse and 169 normalized expression values for human profiles.

### Differential expression, up- and down-regulation

We classified gene expression profiles for 20,316 human protein-coding genes and 22,133 mouse protein-coding genes according to their shape in two steps. In the first step we estimated the overall down-regulation and up-regulation shape of the particular profile. We sorted each expression profile both in ascending and in descending order, corresponding to the “ideal” up-regulated and down-regulated profiles, that is a profile, in which the expression value in a given time-point is higher (lower) than the expression in the previous time-point (Extended Data Fig. 1b). Then, for each gene, we computed the correlation between the chronologically sorted values (the original profile) and the values sorted in ascending and descending order. If a gene has a “perfect” pattern of up-regulation (down-regulation) the correlation will be 1 (−1). The highest value (of the two) was used to classify a gene as potentially up-regulated or down-regulated. We used the p-value corrected for the total 20,316 human and 22,133 mouse observations. Only profiles with False Discovery Rate (FDR) < 0.01 were actually classified as candidate up-regulated or down-regulated. The remaining profiles were classified as irregular.

In the second step we estimated the fold-changes of candidate up- and down-regulated profiles. We first smoothed the profiles using shape constrained (monotonic increase for up-regulated and decrease for down-regulated) additive models (*42*). This allows us to estimate the expression fold-change as a difference between maximum and minimum observed expression values. Since we use log2 expression transformation with one value pseudocount the one unit difference in this scale roughly corresponds to the two-fold difference in the TPM scale. We finally classified the genes with at least one unit difference in log2 expression as up- or down-regulated genes.

### Pace of up- and down-regulation

We used the smoothed gene expression profiles to estimate the speed of up- or down-regulation. We used one unit in the log2 expression scale as a threshold for two-fold up- and down-regulation and estimated the time at which the profile advanced this distance from the 0h time-point value (Extended Data Fig. 1c). Since all the up- and down-regulated genes have at least one unit fold-change difference in log2 scale by construction the estimated pace cannot exceed the total length of time-series, 168 and 72 hours in human and mouse respectively.

### GO analysis

To perform function analysis for a particular group of genes we developed a list of human Ensembl gene identifiers and sent it to Metascape (*43*) (http://metascape.org). For each analysis we plot the resulting “HeatmapSelectedGO.png” files.

### Multiple sequence alignment of CEBPA binding upstream of *MYC* FUSE enhancer

To align the CEBPA binding sites upstream of *MYC* FUSE enhancer we used a multiple sequence alignment of 120 mammals (*44*). This alignment uses either human hg38 or mouse mm10 genomes as reference and builds the alignment of the remaining genomes with respect to them. As a result one nucleotide insertion in all non-reference genomes are ignored in the final alignments (per-reference-chromosome MAF files). To recover these insertions we extracted the alignment that corresponds to our region of interest, selected each individual sequence and aligned it to the corresponding genome using BLAT (*45*) (i.e. the human sequence to the human genome, the rabbit sequence to the rabbit genome, etc). We re-aligned the sequences resulting from querying the genome this way using T-Coffee (*46*).

### Processing of ChIP-Seq data

For the ChIP-Seq experiments published elsewhere we downloaded corresponding raw read sequences from the GEO database (*47*). The ChIP-Seq sequence reads were aligned to human (hg38) and mouse (mm10) genomes using BWA (*48*), for compatibility we merged replicated experiments wherever possible. We removed potential duplicated reads with Picard (*49*). We called peaks comparing ChIP (immunoprecipitation) signal with corresponding input control. Since mouse CEBPA ChIP-Seq data along transdifferentiation of C10 cells (*4*) do not have associated input data we used the input from mouse B cells (*7*).

To call the peaks we used GEM peak caller (*50*) providing the estimated mappable genome size of 2,424,833,933 nucleotides for human and 2,107,021,337 nucleotides for mouse genomes. We run GEM in pure peak calling mode without identifying binding motifs and added “--relax” parameter in order to get more peaks. We used MACS2 (*51*) to develop genome-wide fold-change and SPMR (signal per million reads) values. Running MACS2 we set the mappable genome size parameters equal to GEM ones, set the fragment length parameter equal to 250 for all runs and demanded MACS2 to normalize reads counts with “--SPMR --bdg” parameters. In total we generated three different genome-wide signal tracks (bigWig files). The first track corresponds to immunoprecipitation (SPMR or normalized pileup) signal, the second track corresponds to fold over input (fold-change) signal. To properly generate corresponding input/control SPMR signal values we ran MACS2 using the alignment of input reads as fake immunoprecipitation data and stored the corresponding SPMR track as a true input/control signal. This procedure allows us to correctly compare the levels of CEBPA binding in repeated regions (Supplementary Table 2).

Within the whole manuscript reporting the results of our ChIP-Seq analysis we show the peaks counts and positions of each peak following the GEM peak caller. We report fold-change and SPMR signals as maximum values within corresponding regions following MACS2 peak caller. Reporting the images from UCSC genome browser (*52*) we plot corresponding fold-change bigWig files.

### ChIP Atlas analysis

The ChIP Atlas database (*18*) collects information about ChIP-Seq experiments archived in the NCBI Sequence Read Archive (SRA) (*53*). Peaks are called using an uniform pipeline that runs MACS2 only on immunoprecipitation data, without using any input/control. Peaks are called at different FDR thresholds (10^-5^, 10^-10^, 10^-20^), and normalized immunoprecipitation signals as SPMR bigWig files are reported. We used the peaks at the 10^-5^ FDR threshold and the corresponding bigWig files. When reporting ChIP Atlas results data, we consider the maximum SPMR values within the peaks.

### Reproducible CEBPA peaks

Based on the optimal correspondence of human and mouse time-points (Supplementary Methods), we selected six human (3h, 6h, 12h, 48h, 72h and 120h) and six mouse (30m, 1h, 3h, 12h, 24h and 48h) CEBPA ChIP-Seq experiments. We merged all human (mouse) peaks and selected the regions that were called peaks in at least three time-points as reproducible human (mouse) peaks. To calculate average genome peak density we used mappable genome size for human and mouse genomes calculated by GEM above.

### Candidate targets genes

We used human Gencode v30 and mouse Gencode vM22 gene and transcript annotations. Gencode annotates one of the transcripts as a major isoform. We used the transcription start site of this isoform as an anchor for every gene. In the case of MYC we selected the (−2,000: +500) region around this anchor and in the case of CEBPA the region (−25,000: +500). We overlapped these regions with MYC ChIP-Seq peaks, and with reproducible CEBPA peaks. We called a gene, a MYC or CEBPA candidate target if the region contained a MYC or reproducible CEBPA peak.

### CEBPA binding sites within human and mouse genomes

We downloaded the masked versions of hg38 and mm10 genomes from the UCSC browser. We scanned masked and unmasked genomes with either the human CEBPA_HUMAN.H11MO.0.A (Fig. 3) or the mouse CEBPA_MOUSE.H11MO.0.A (Extended Data Fig. 5) position weight matrices (PWMs) from the Hocomoco database (*14*) using SPRy-SARUS motif scanner (*54*). We set a minimum threshold for motif score equal to zero and selected the binding site with the highest score from overlapping predictions in the different strands since the CEBPA motif is highly palindromic. We estimated the distribution of motif scores using the Kernel Smoothing R package (*55*), binning the scores (from 0 to 10,3) into 10,301 bins with band-width equal to 0.00538 for human and 0.00485 for mouse CEBPA PWMs. We calculated the enrichment of CEBPA motifs in human repetitive regions dividing the number of matches of the CEBPA motif with corresponding score in the unmasked genome by the number of matches in the masked genome.

### CEBPA binding in human repetitive regions

We selected seven CEBPA motifs with PWM scores greater than 7 and repeat enrichments greater than 6 for subsequent analysis. We downloaded the annotations of all human repetitive regions from UCSC. To calculate the type of repetitive elements corresponding to the enriched binding motifs, we overlap the DNA positions of the corresponding binding motifs instances with this annotation. We found the motifs corresponding to the enriched scores to map almost exclusively to Alu repeats, with some mapping also to L1 and Satellite repeats (Supplementary Table 2). To investigate CEBPA ChIP-Seq binding in human repetitive regions we first extract DNA sequences for all Alu, L1 and Satellite repeats within the human genome and scan them for CEBPA binding sites with CEBPA_HUMAN.H11MO.0.A matrix. We selected 431,323 Alu, 192,636 L1 and 1,387 Satellite regions that contain a CEBPA binding motif with score above 7 for subsequent analysis. We selected the maximum SPMR within each repeat for both the CEBPA ChIP-Seq and input signal bigWig files (Supplementary Table 2 for the genome wide average at each transdifferentiation time-point). We used a t-test to test the significance of the enrichment of the immunoprecipitation compared with the input signal.

### Analysis of Hi-C data

We downloaded Hi-C data in the THP1 cell line from ENCODE portal (*56*), accession ENCSR748LQF. We merged all the replicates. Our analysis followed the original Juicer pipeline (*57*). We run the Juicer scripts manually: BWA (*48*) for read alignments, “chimeric_blacklist.awk” to scan for properly aligned reads pairs, “fragment.pl” to annotate fragments, then manually sorted read pair alignments and removed duplicated ones. We finally run the “juicer_tools.jar pre” program to create the Hi-C contact matrix. We used Juicebox (*58*) to visualize the Hi-C contacts for the human *ID2* gene at 5 kb resolution (Extended Data Fig. 8).

## ACKNOWLEDGMENTS

We thank Francesca Rapino and Thomas Graf for advice on the transdifferentiation experiments, Alessandra Breschi for assistance with RNA-Seq data analysis and Marina Ruiz Romero for advice on transgene experiments. We also thank the Genomics and the Flow Cytometry Units of the CRG (Barcelona, Spain), for sample processing. Special thanks to Romina Garrido for administrative support.

## Funding

ERC’s/European Community under the FP7 (ERC-2011-AdG-294653-RNA-MAPS)

Spanish Ministry of Economy and Competitiveness (MEC) (BIO2011-26205).

Spanish Ministry of Science and Innovation to the EMBL partnership

Centro de Excelencia Severo Ochoa from the CERCA Programme / Generalitat de Catalunya.

This work reflects only the author’s views and that the Community is not liable for any use that may be made of the information contained therein.

## Author contributions

R.N. analyzed the data, S.P.L. and C.A. carried out transgene experiments, M.S., A.A. and A.E. generated RNA-Seq and ChIP-Seq data, S.U. contributed to data analyses, S.P.L. and R.J. have coordinated wet-lab experiments, R.G. conceived and supervised the project. R.N., S.P.L. and R.G. wrote the manuscript.

## Data and materials availability

RNA-Seq and ChIP-Seq reads generated during the current study were deposited in the ArrayExpress database at EMBL-EBI (www.ebi.ac.uk/arrayexpress) under accession numbers E-MTAB-7115 and E-MTAB-7114 respectively. All data are available in the main text or the supplementary materials.

## Code availability

All algorithms used for the data analysis are described in the Methods and Supplementary Methods sections.

**Supplementary Information** is available for this paper.

Correspondence and requests for materials should be addressed to Dr Roderic Guigo for general questions and regarding publication process, to Dr Silvia Perez-Lluch regarding wet-lab procedures and to Dr Ramil Nurtdinov for dry-lab procedures and improvement or reorganization of the figures.

---

**Extended Data Fig. 1.**
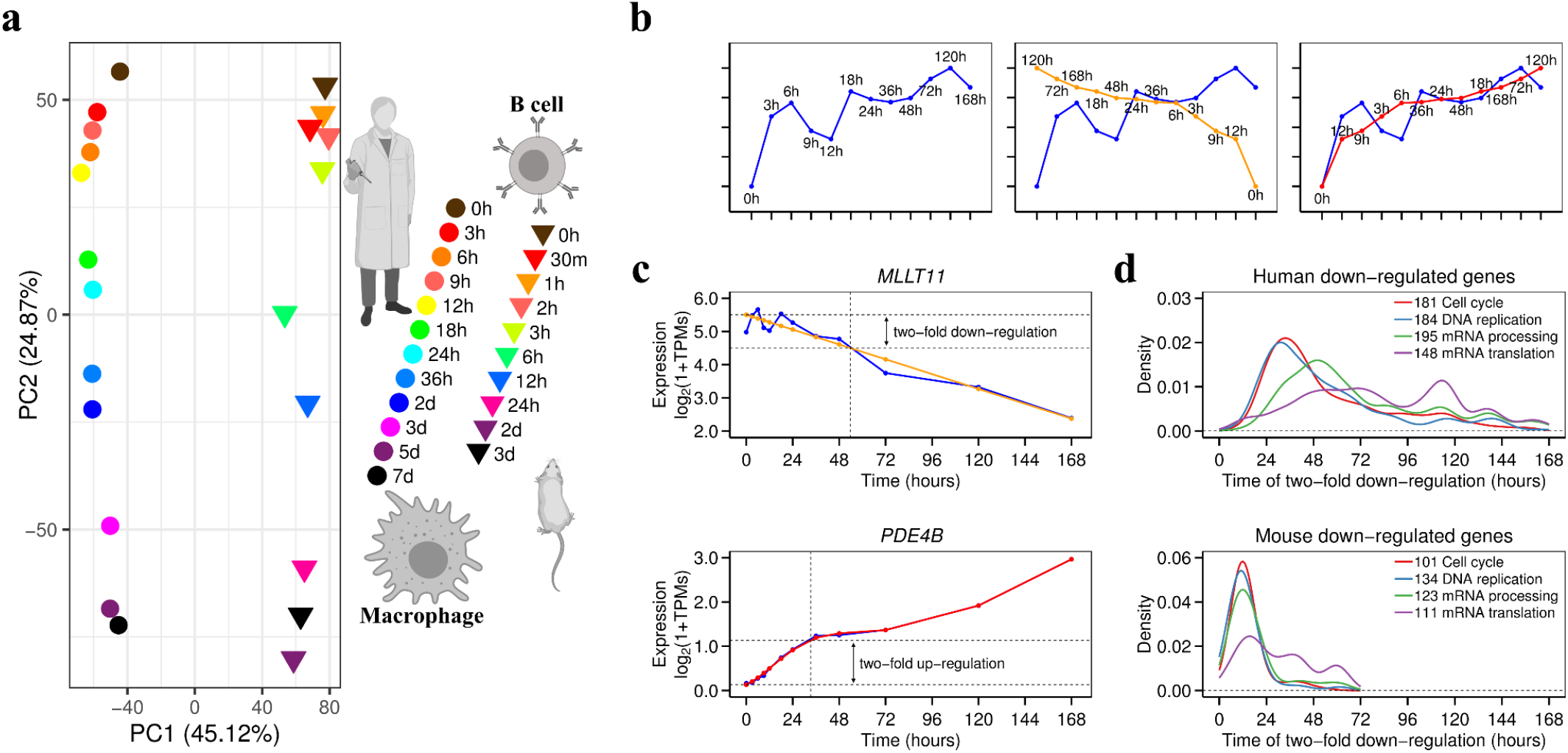
Transcriptional dynamics during transdifferentiation. **a,** PCA of the transdifferentiation samples in human and mouse based on the expression of 9,188 human-mouse expressed orthologous genes. **b**, Identification of up-regulated (down-regulated) genes. Gene expression profile of the human *CCNL1* gene (left panel). Comparison of the real and “ideal” down-regulated profile (middle panel) and “ideal” up-regulated profile (right panel) for *CCNL1*. The correlation of the real with the up-regulated profile is 0.840, and with the down-regulated profile is −0.663. We assumed the gene to be up-regulated. To consider a gene significatively up- or down-regulated we used a threshold of FDR < 0.01. In this case, FDR=0.000775, therefore the gene *CCNL1* is considered up-regulated in human BLaER1 cells. **c,** Time of two-fold expression change. Top panel - the time of two-fold down-regulation for MLLT11 is 53.7 hours (the projected time at which the smoothed expression of the gene is two-fold down the expression at the start of the transdifferentiation). Bottom panel - the time of two-fold up-regulation for PDE4B is 33.3 hours (the projected time at which the smoothed expression of the gene is two-fold up the expression at the start of the transdifferentiation). Since we are using a logarithmic scale for expression, two-fold expression change corresponds to one unit in this scale. **d,** Distributions of two-fold down-regulation times for human (top) and mouse (bottom) genes related to cell division and RNA metabolism.

**Extended Data Fig. 2.**
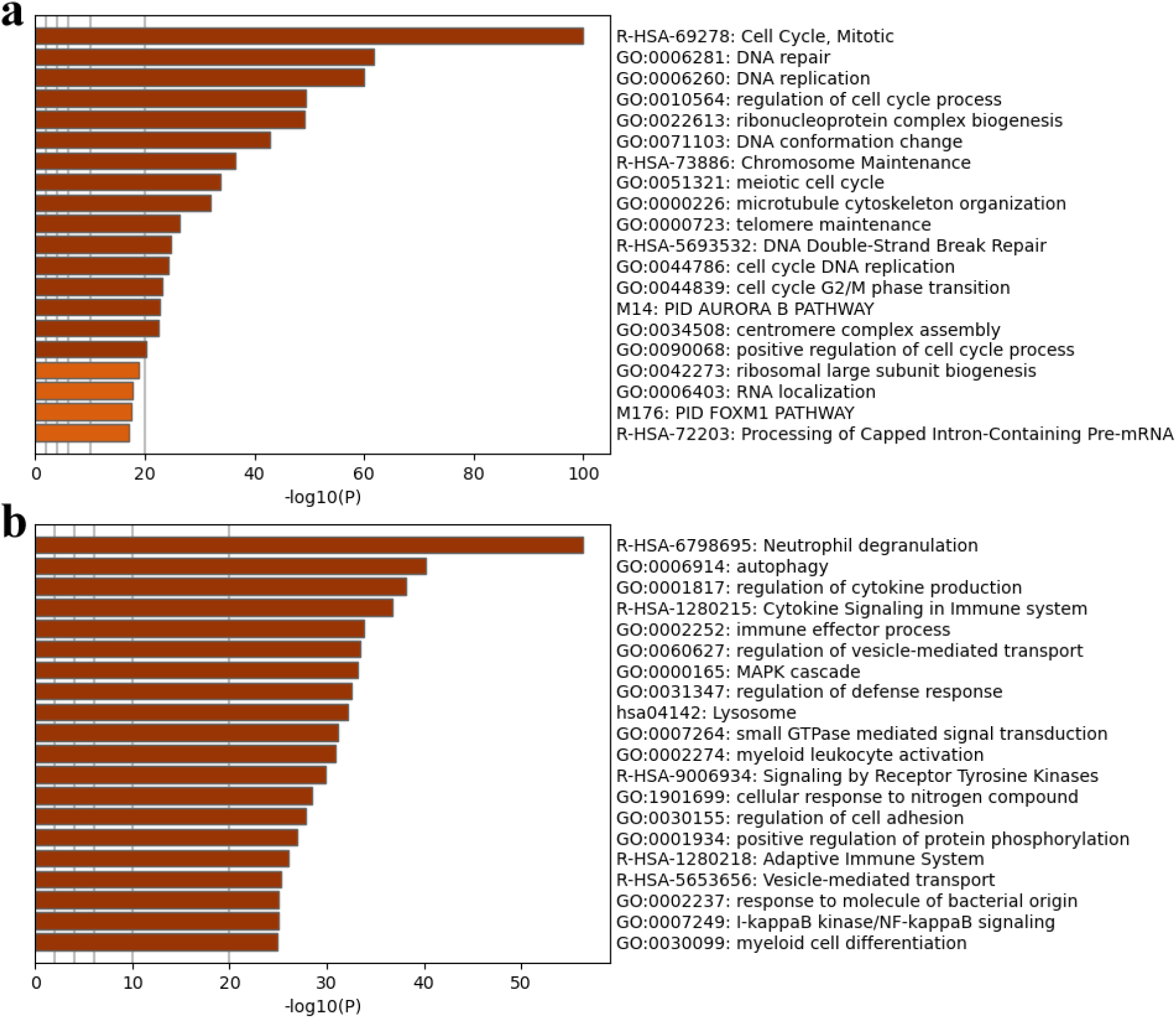
Functional annotation of up- and down-regulated genes using Metascape (***43***). **a,** Functional enrichment of 614 genes down-regulated during transdifferentiation of both human BLaER1 and mouse C10 cells. **b,** Functional enrichment of 1,317 genes up-regulated during transdifferentiation of both human BLaER1 and mouse C10 cells.

**Extended Data Fig. 3.**
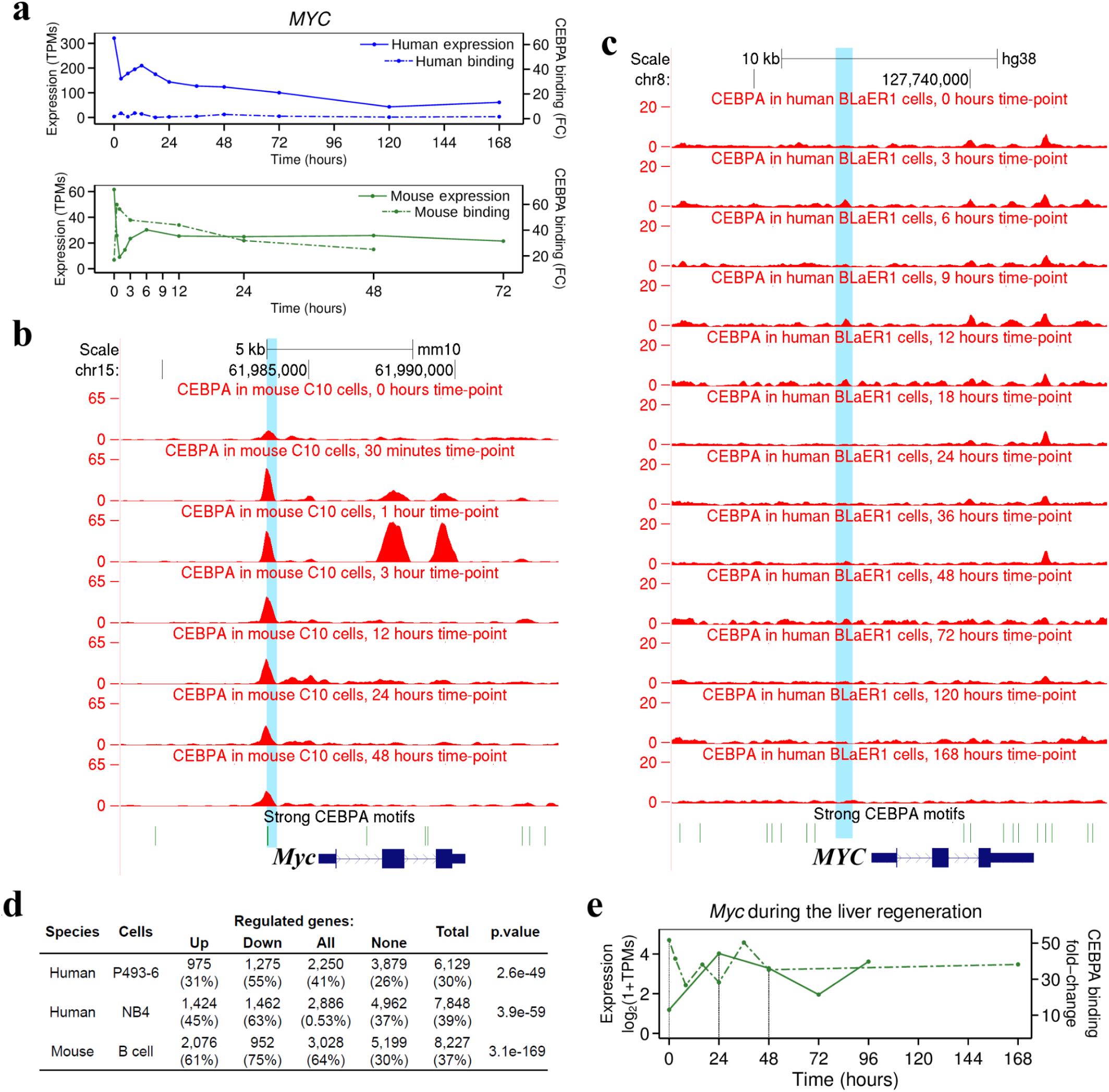
*MYC* expression and CEBPA binding at the *MYC* locus. **a,** Complete expression profiles for human (top panel) and mouse (bottom panel) *MYC* expression and CEBPA binding at the FUSE enhancer. **b,** Fold-change signal of CEBPA binding at mouse *Myc* locus at seven time-points during transdifferentiation of C10 cells. **c,** Fold-change signal of CEBPA binding at *MYC* locus at twelve time-points during transdifferentiation of BLaER1 cells. **d,** MYC candidate target genes in human and mouse. P.value corresponds to the Chi-Squared test comparing Regulated and Non-regulated genes. Numbers within brackets correspond to the percentage of the candidate target genes among all genes within the corresponding groups. **e,** Dynamics of CEBPA binding and *Myc* expression at the *Myc* FUSE during liver regeneration. Unaffected liver shows strong CEBPA binding, and very low levels of *Myc* (0h). The regenerating liver, after partial hepatectomy, shows decreased CEBPA binding and increased *Myc* expression. Since not all hepatocytes participate in regeneration, the bulk omics data is likely to attenuate the actual magnitude of the dynamics. Overall, along four different time-points, we found, as during transdifferentiation, strong negative correlation between CEBPA binding and Myc expression (cc=-0.95, p-val=0.0469).

**Extended Data Fig. 4.**
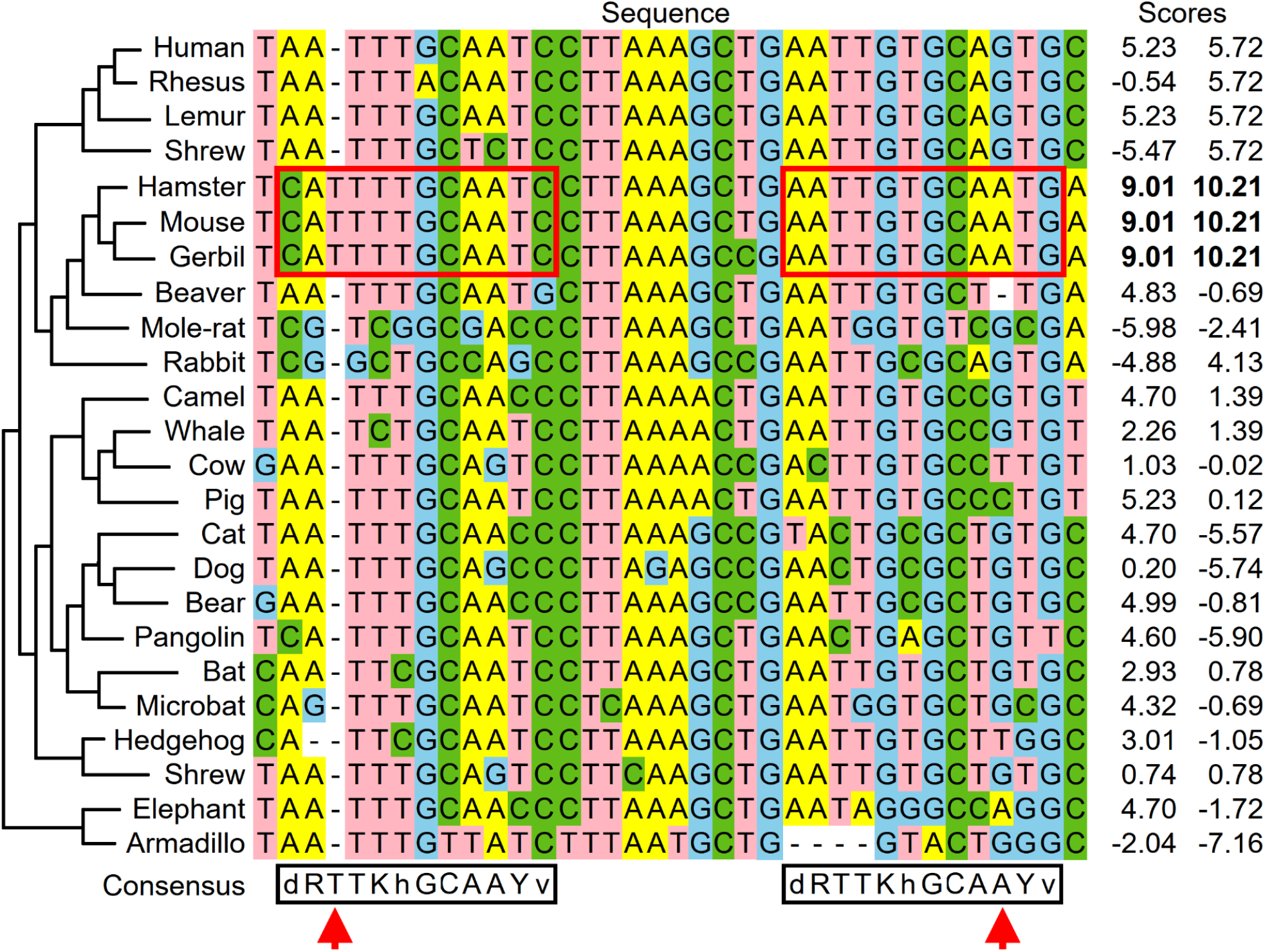
CEBPA binding sites in *MYC* FUSE enhancer. Multiple sequence alignment, across selected mammals, of the region within the FUSE enhancer containing two very strong CEBPA binding sites in muroidea. These sites are enclosed in red boxes in the figure. Positions in which mutations confer higher affinity for CEBPA are marked with an arrow. The displayed human and mouse sequences start 1,813 and 1,759 bp, respectively upstream from the *MYC* TSS. The scores of the binding sites (right) are computed using the human CEBPA PWM from Hocomoco (*14*). The Hocomoco CEBPA binding consensus is shown at the bottom of the alignment.

**Extended Data Fig. 5.**
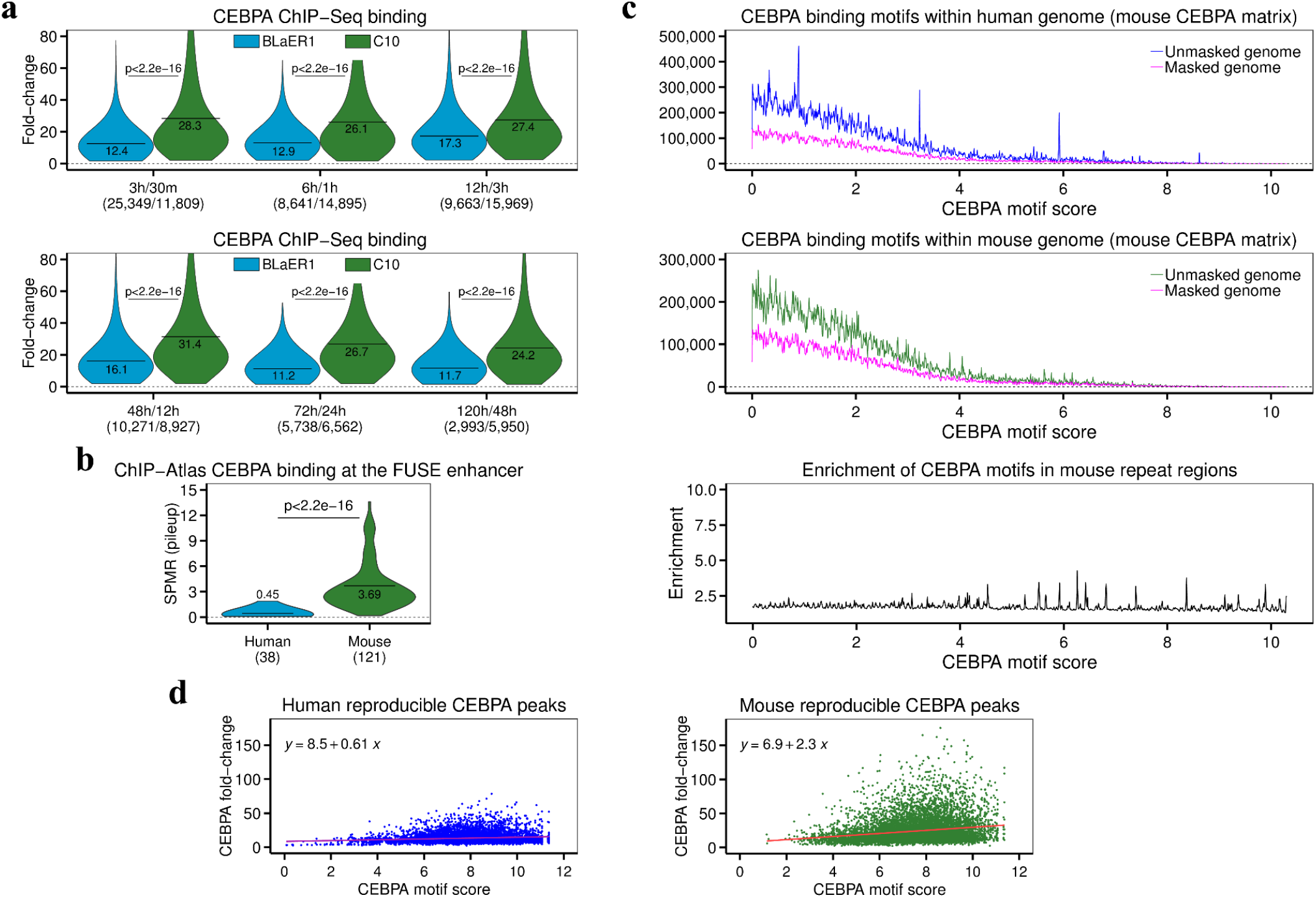
CEBPA binding in human and mouse. **a,** Distribution of fold-change of CEBPA peaks in the human and mouse genome at matched time-points. **b**, Strength of CEBPA binding (SPMR, Signal Per Million Reads) at the human and mouse *MYC* FUSE enhancer (in parenthesis number of CEBPA ChIP-Seq experiments in ChIP-Atlas). In **a** and **b**, horizontal lines represent the mean value. **c,** Distribution of the strength of CEBPA binding sites within the human genome (top panel). Enrichment of CEBPA binding sites in mouse repetitive regions (middle and bottom panel). For each PWM score, the enrichment is the number of matches of the CEBPA motif with this score in the unmasked genome over the number of matches in the masked genome. The enrichment is very poor compared to that observed in human (Fig. 3d). **d,** CEBPA binding fold-change versus score of the underlying CEBPA binding motif for 8,494 reproducible human peaks (left panel). CEBPA binding fold-change versus score of the underlying CEBPA binding motif for 10,115 reproducible mouse peaks (right panel). In the upper-left corner of the corresponding plots the parameters of the linear regression are displayed.

**Extended Data Fig. 6.**
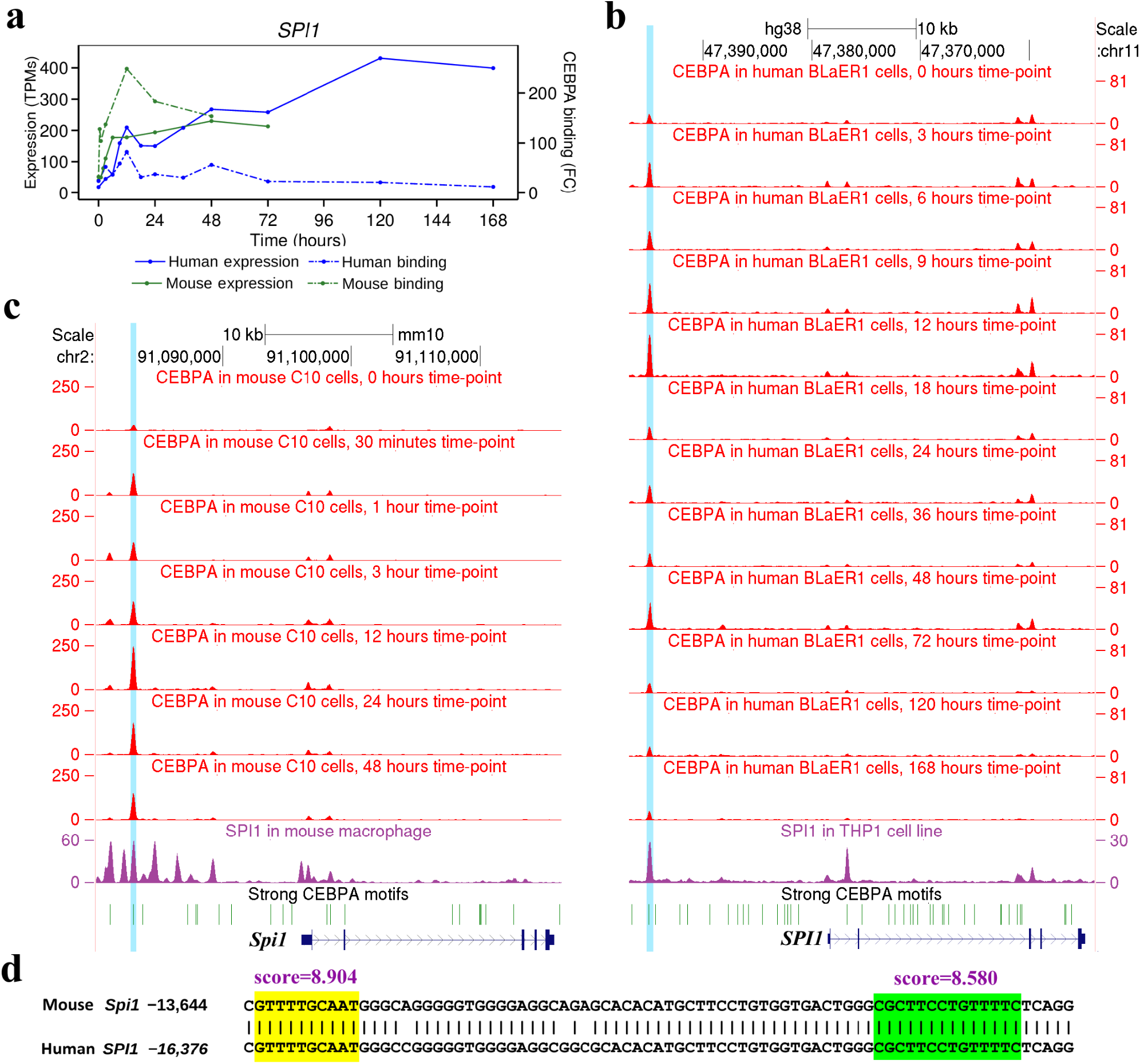
*SPI1* expression and CEBPA binding at the *SPI1* locus. **a,** Expression profiles for human and mouse *SPI1* expression and CEBPA binding at the *SPI1* upstream enhancer. **b,** Fold-change signal of CEBPA binding at *SPI1* locus at twelve time-points during transdifferentiation of BLaER1 cells, and of SPI1 binding in human THP1 cells (*16*). **c,** Fold-change signal of CEBPA binding at the mouse *Spi1* locus at seven time-points during transdifferentiation of C10 cells, and of SPI1 binding in mouse macrophages (*59*). **d,** Human-mouse DNA alignment of the upstream enhancers of *SPI1* gene. Positions in the alignment are indicated with respect to *SPI1* TSS. The yellow box indicates a predicted CEBPA binding site, the green box indicates a predicted SPI1 binding site. Such co-localization of SPI1 and CEBPA binding has been shown to up-regulate many myeloid-specific genes (*60*).

**Extended Data Fig. 7.**
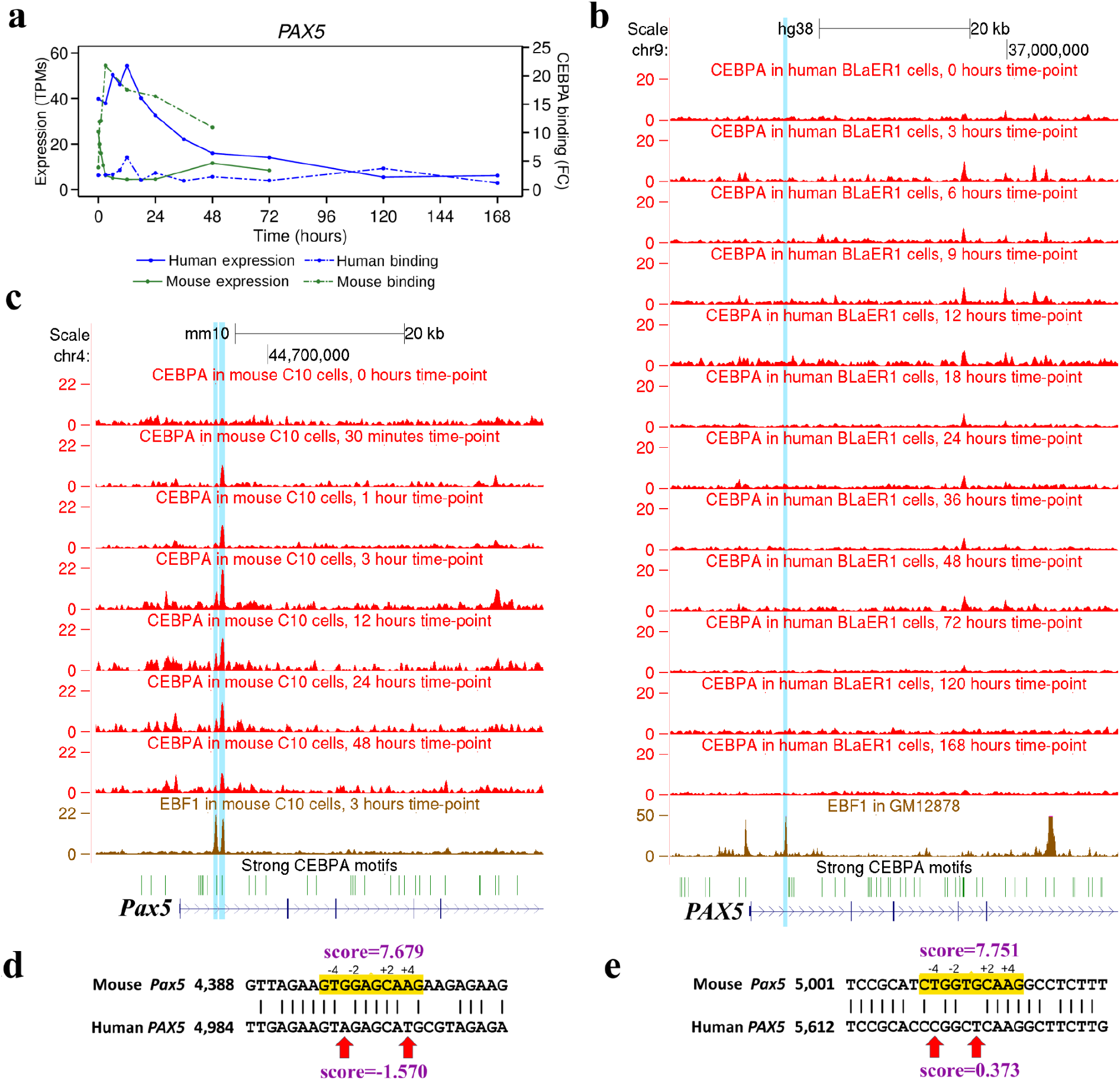
*PAX5* expression and CEBPA binding at the *PAX5* locus. **a,** Expression profiles for human and mouse *PAX5* expression and CEBPA binding at the first intron enhancer. **b,** Fold-change signal of CEBPA binding at *PAX5* locus at twelve time-points during transdifferentiation of BLaER1 cells and of EBF1 in GM12878 cells (*61*). **c,** Fold-change signal of CEBPA and EBF1 binding at mouse *Pax5* locus during transdifferentiation of C10 cells. CEBPA binding is for seven time-points and EBF1 for 3 hours after induction (*4*). **d-e,** Human-mouse DNA alignment of the CEBPA binding motifs within the proximal and distal first intron regulatory enhancers of the *PAX5* gene. Positions in the alignment are indicated with respect to *PAX5* TSS. The yellow box indicates a predicted CEBPA binding site, red vertical arrows indicate the substitutions that substantially alter the motif strength between human and mouse.

**Extended Data Fig. 8.**
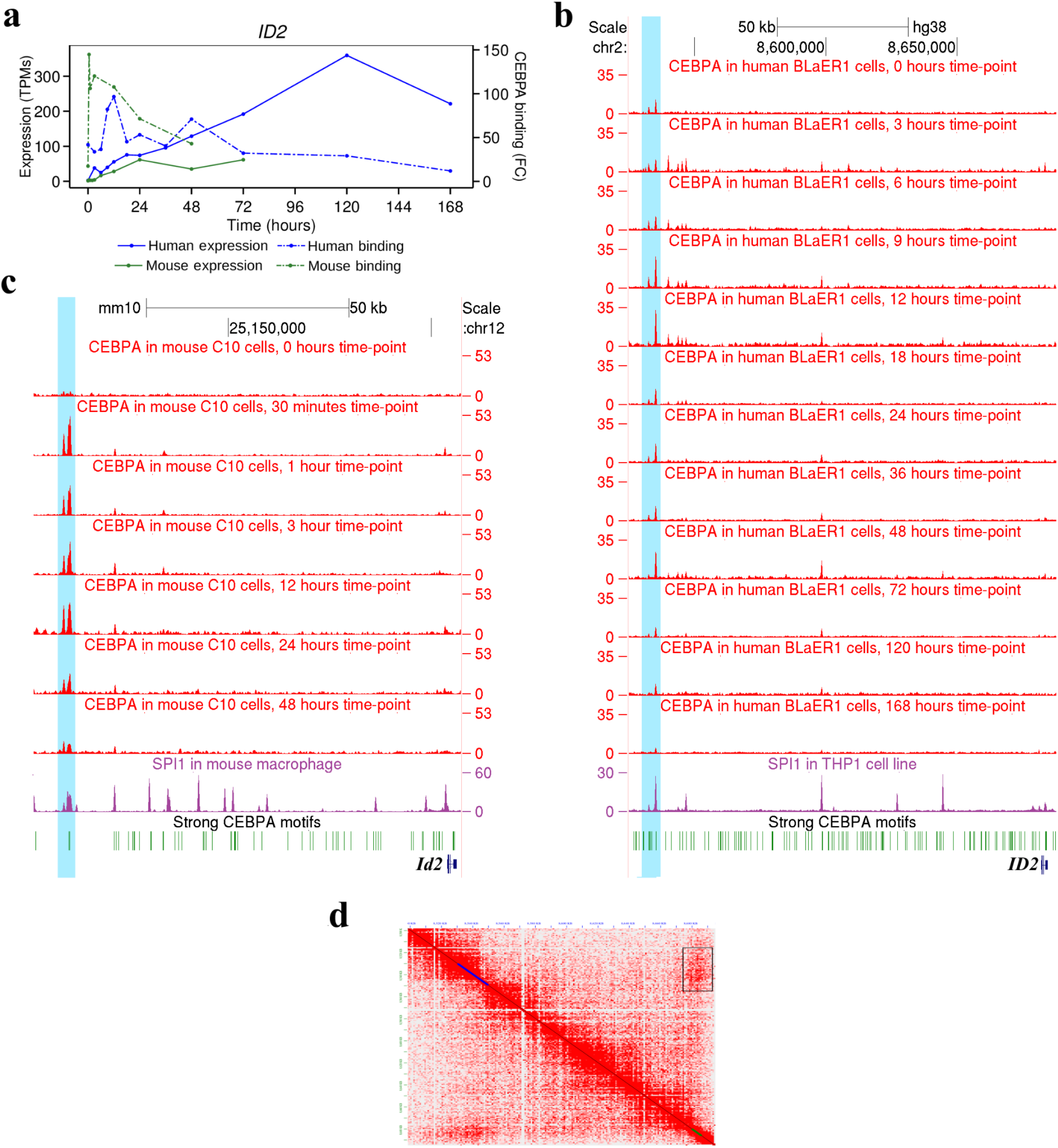
*ID2* expression and CEBPA binding at the *ID2* locus. **a,** Complete expression profiles for human and mouse *ID2* expression and CEBPA binding in a predicted distal (150 kb in human and 80 kb in mouse) upstream enhancer. **b,** Fold-change signal of CEBPA binding at *ID2* locus at twelve time-points during transdifferentiation of BLaER1 cells, and of SPI1 binding in human THP1 cells (*16*). **c,** Fold-change signal of CEBPA binding at mouse *Id2* locus at seven time-points during transdifferentiation of C10 cells, and of SPI1 binding in mouse macrophages (*59*). **d,** Chromosome conformation capture Hi-C data at *ID2* locus in the human THP1 cell line (*56*). The region corresponding to the ID2 gene body is highlighted in green. This region shows enriched contact density (black rectangle in the upper right corner) with a region 149 kbp upstream (highlighted in blue). The conserved CEBPA binding between human and mouse in panels A-C occurs in this predicted enhancer region.

**Extended Data Fig. 9.**
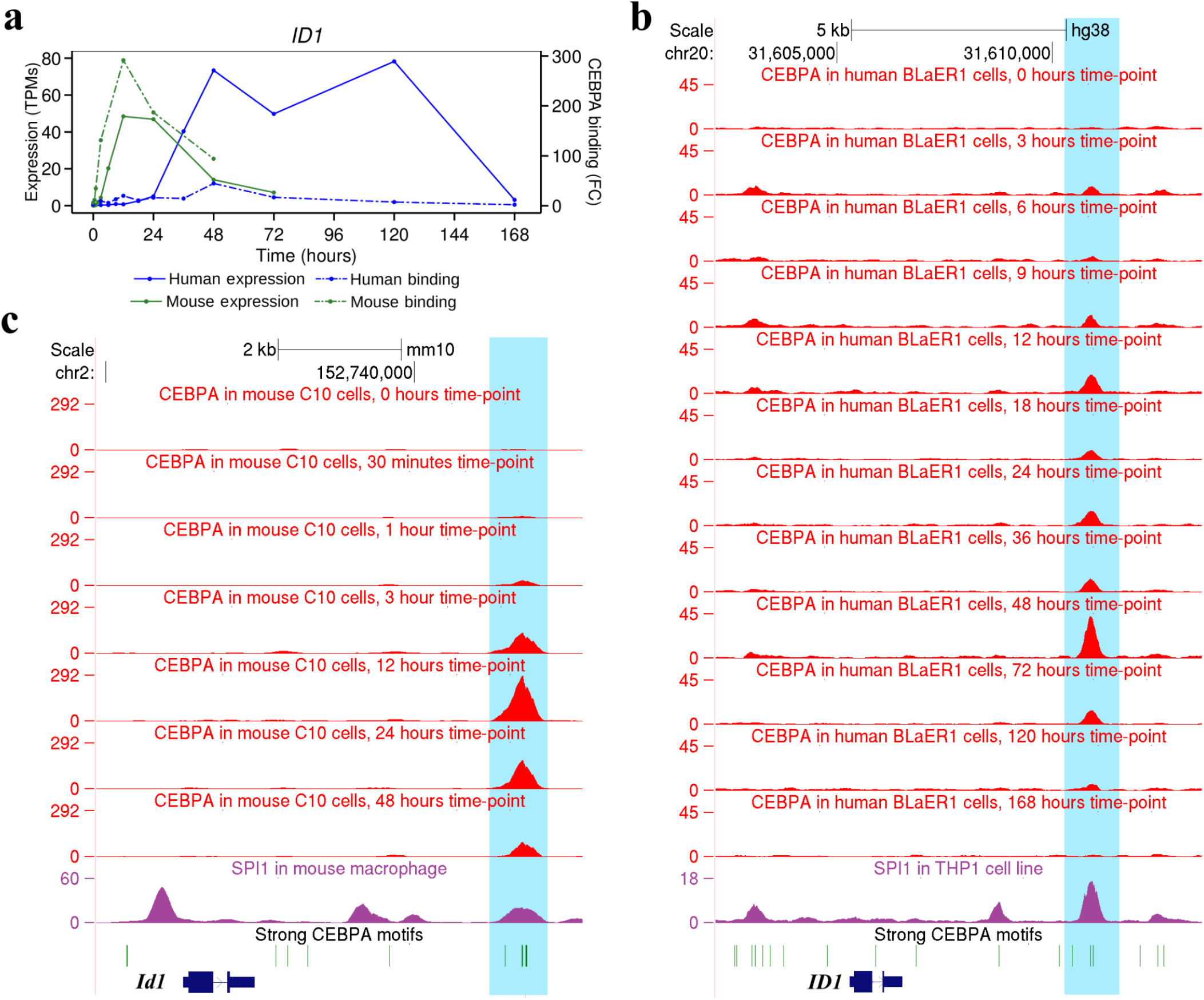
*ID1* expression and CEBPA binding at the *ID1* locus. **a,** Complete expression profiles for human and mouse *ID1* expression and CEBPA binding in a potential downstream enhancer. **b,** Fold-change signal of CEBPA binding at *ID1* locus at twelve time-points during transdifferentiation of BLaER1 cells, and of SPI1 binding in human THP1 cells (*16*). **c,** Fold-change signal of CEBPA binding at mouse *Id1* locus at seven time-points during transdifferentiation of C10 cells, and of SPI1 in mouse macrophages (*59*).

**Extended Data Fig. 10.**
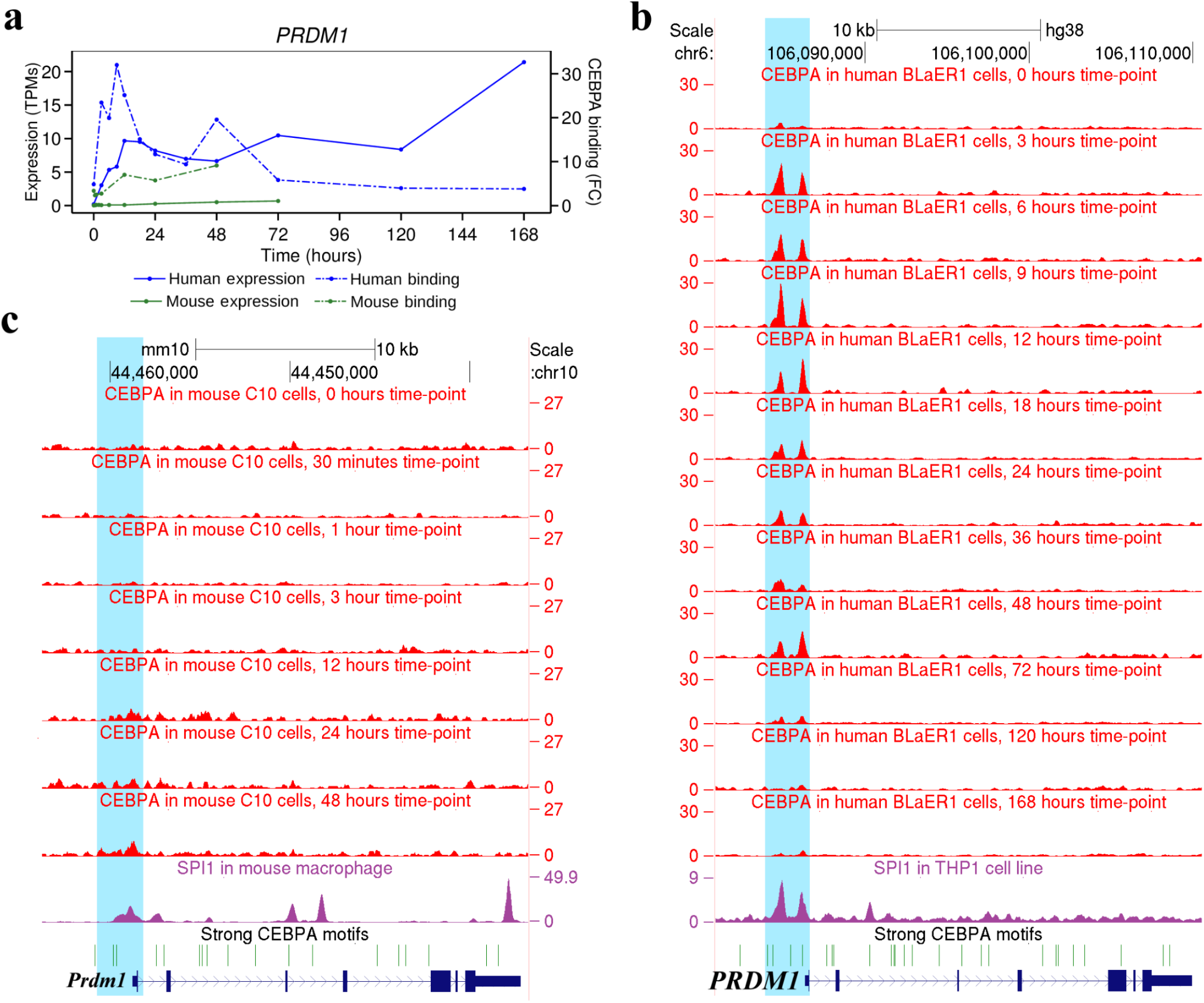
*PRDM1* expression and CEBPA binding at the *PRDM1* locus. **a,** Complete expression profiles for human and mouse *PRDM1* expression and CEBPA binding in its promoter region. **b,** Fold-change signal of CEBPA binding at *PRDM1* locus at twelve time-points during transdifferentiation of BLaER1 cells, and of SPI1 binding in human THP1 cells (*16*). **c,** Fold-change signal of CEBPA binding at mouse *Prdm1* locus at seven time-points during transdifferentiation of C10 cells, and of SPI1 in mouse macrophages (*59*).

## Supplementary Methods

### Cloning of reporter plasmids

For intermediate reporter plasmid, the *mCherry* gene was PCR amplified from plasmid pDECKO_mCherry (Addgene 78534) with primers mCherry_Fw and mCherry_Rv using Expand polymerase (Roche). The PCR product was inserted into a lentiGuide-Puro plasmid (Addgene 52963) digested with *KlfI* (Thermo Fisher) and *BsmbI* (Thermo Fisher) by Gibson cloning (*62*). Constructs were transformed into z-Stbl3 competent cells. Genomic DNA from human and mouse cells was extracted with the GeneJet Genomic purification kit (Thermo Scientific). The human *MYC* promoter was amplified with primers hMYC_Fw and hMYC_Rv and KOD polymerase (Takara). The mouse *Myc* promoter was amplified with primers mMyc_Fw and mMyc_Rv and KOD polymerase. Human and mouse promoters were inserted into LentiGuide-Puro-intermediate-mCherry-reporter digested with *KlfI*.

For cloning of human and mouse plasmids with mutations in the enhancer regions, we use the non-mutated plasmids as a template for 2 PCR reactions, obtaining 2 different inserts that were cloned together with intermediate plasmid (LentiGuide-Puro-intermediate-mCherry-reporter), previously digested with *KlfI*. The human *MYC* mutant enhancer was obtained by amplification of plasmid LentiGuide-Puro-hMyc-mCherry using primers sets: hMYC_Fw and hMYC_2_Rv for insert-1, and hMYC_2_Fw and hMYC_Rv for insert-2. The mouse *Myc* mutant enhancer was obtained by amplification of plasmid LentiGuide-Puro-mMyc-mCherry using primers sets: mMyc_Fw and mMyc_2_Rv for insert-1, and mMyc_2_Fw and mMyc_Rv for insert-2. Positive clones were tested by colony PCR and Sanger sequencing using primers: pLenti_seq_Fw, pLenti_seq_Rv, internal_hMYC_Fw (for sequencing internal *hMYC*) and internal_mMyc_Fw (for sequencing internal *mMyc*). The primer sequences used for cloning and Sanger sequencing can be found in Table S7.

### Cell transfection and infection

For lentivirus production, HEK293T virus packaging cells, cultured in standard supplemented medium, were co-transfected with each pLenty-mCherry construct, pVsVg packaging plasmid (Addgene 8484) and psPAX2 vector (Addgene 12260) using Lipofectamine 2000 (according to the manufacturer’s protocol). Viral supernatant was collected 48h post transfection and used for spin infection of 1.5E106 BLaER1 and C10 cells at a density of 500,000 cells/ml for 3 hours at 1,000G at 30°C. The percentage of infection was checked with a Fortesa cell cytometer analyser. After 48h of infection, the cells were double selected with puromycin (2 μg/ml) and blasticidin (20 μg/ml) for two weeks. The cells were further selected by Fluorescence-Activated Cell Sorting in a BD Influx sorter before induction of transdifferentiation. Induction of transdifferentiation was performed as previously described (*1, 2*). Cells were collected at times 0, 1, 3 and 6 hours after induction and split for chromatin immunoprecipitation and expression analysis of reporter experiments.

### Expression profile alignment

We refer to the vector of 12 (human) or 10 (mouse) expression values for a given gene along transdifferentiation as the expression profile of this gene. We aligned human and mouse expression profiles of orthologous genes using dynamic time warping algorithm (*63*). For a given orthologous gene, we first z-normalized the human and mouse profiles independently and aligned them. As a distance D_i,j_ between i-th human and j-th mouse time-points we used the absolute value of their z-normalized expression difference [1]. Following the standard dynamic programming algorithm we then filled recursively the distance matrix M_i,j_ [2-5]:

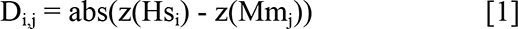

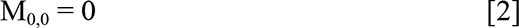

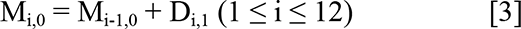

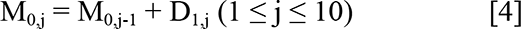

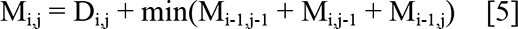

The bottom right value of the cumulative matrix, M_12,10_ corresponds to the minimum distance between profiles. To infer the optimal alignment we trace back the matrix starting from M_12,10_. At each comeback step we selected the cell with lowest distance until we reached M_0,0_.

To decide whether the distance of the aligned profiles was significantly smaller than the one of a random alignment, we computed the background distribution of the alignment distances, permuting the gene expression profiles 250,000 times per gene. Within each permutation we calculated the minimal distance between shuffled profiles, and estimated as a p-value the proportion of random alignments in which the distance was smaller than alignment distance of the actual profiles. We used an FDR ≤ 0.05 correcting for the 9,188 genes to consider the aligned profiles more similar than expected by chance. In this case, we refer to the human and mouse profiles as “concordant”. Otherwise, we refer to them as “discordant”.

Based on profile alignments of 2,726 concordant genes we obtained the optimal correspondence between human and mouse time-points. To each time-point in mouse, we assigned the time-point in human with the maximum number of concordant alignments. The optimal correspondence assigns time 0h in human and 0h in mouse (2,001 alignments), 3h in human and 30m in mouse (1,237 alignments), 6h and 1h (793 alignments), 9h and 2h (659 alignments), 12h and 3h (585 alignments), 18h and 6h (546 alignments), 48h and 12h (801 alignments), 72h and 24h (1,270 alignments), 120h and 48h (1,616 alignments) and 168h and 72h (2,109 alignments).

### Down Sampling of ChIP-Seq data

We performed CEBPA ChIP-Seq data for 12 human time-points in two replicates. On average among replicates we sequenced 43 million immunoprecipitation reads and 168 million input reads per time-point. Previously published mouse CEBPA ChIP-Seq data for seven time-points have on average only 15.6 million reads per time-points (Supplementary Table 1). Sequence depth can influence both the number and strength of the peaks. The more the reads sequenced, the higher the chance to identify weaker peaks, which, in turn, decreases the peak fold-change. To control the impact that the difference in sequencing depth may have in the differences in binding strength that we observed between human and mouse, we downsampled the human reads to the average number of mouse reads at each time-point. We used Picard to randomly select on average 15.62 million reads from our human CEBPA ChIP-seq data and re-processed the resulting reads to call peaks as described above. The average fold-change in mouse is 25.785. The average fold-change in human before downsampling is 12.997. After downsampling it becomes 15.410. Thus, as expected, the fold-change in human becomes higher after downsampling, but differences in sequence depth explain only a small fraction of the differences in fold-change observed between human and mouse.

### Reassignment of reads in Alu repeats

We found Alu repetitive regions with strong CEBPA binding sites attracting an excess of CEBPA ChIP-Seq reads compared to input. To assess whether this could explain the difference in CEBPA binding strength observed between human and mouse, we attempted to simulate CEBPA binding if Alu repeats would not attract this excess of reads. We do not pretend this to be a rigorous simulation, which is extremely difficult. The aim of this analysis is simply to obtain a lower bound estimate of the decrease in average CEBPA peak strength (fold-change) that can be explained by the excess of CEBPA reads in Alu repeats. We follow a quite conservative approach in which we simply re-distributed CEBPA ChIP-Seq reads mapping to peaks overlapping Alu repeats, containing strong CEBPA binding sites, to CEBPA peaks not overlapping Alu repeats.

More in detail, for each human time-point we overlapped CEBPA peaks and Alu regions and separated peaks into three groups:

1. Non-Alu peaks that either do not overlap Alu repeats or overlap Alu containing weak (score < 3) CEBPA binding motifs.

2. Intermediate Alu (Int-Alu) peaks that overlap Alu containing intermediate strength (3 ≤ score < 7) CEBPA binding motifs.

3. Strong-Alu peaks that overlap Alu containing strong (score ≥ 7) CEBPA binding motif. Simulating the CEBPA binding in the Alu-free genome we first separated CEBPA read alignments into the donor and recipient sets. CEBPA recipients correspond to read alignments in Non-Alu peaks. CEBPA donors correspond to read alignments in Strong-Alu peaks. Then, we replaced every read from the donor set with a copy of a random read from the recipient set. This procedure allowed us to respect the strength of the Non-Alu peaks while redistributing the reads. We next run GEM and MACS2 peak callers as with the original ChIP-Seq datasets, with exception of the options that switch off the filtering of duplicates (“--nrf” option for GEM and “--keep-dup all” option for MACS2).

Results are presented in Supplementary Table 3. As it is possible to see, after re-distributing peaks from Strong-Alu peaks, some Strong-Alu peaks remain. These Strong-Alu peaks do not overlap the original Strong-Alu peaks. Read redistribution increases the significance of Non-Alu peaks, and, as a consequence a few peaks, which did not pass FDR thresholds originally, become significant afterwards. We also observed a small decrease in the number of Int-Alu peaks. Some Int-Alu peaks lie very close to Strong-Alu peaks, therefore some read alignments may overlap neighboring peaks. Removing the read alignments from Strong-Alu peaks resulted in removing them from neighbor Int-Alu ones as well.

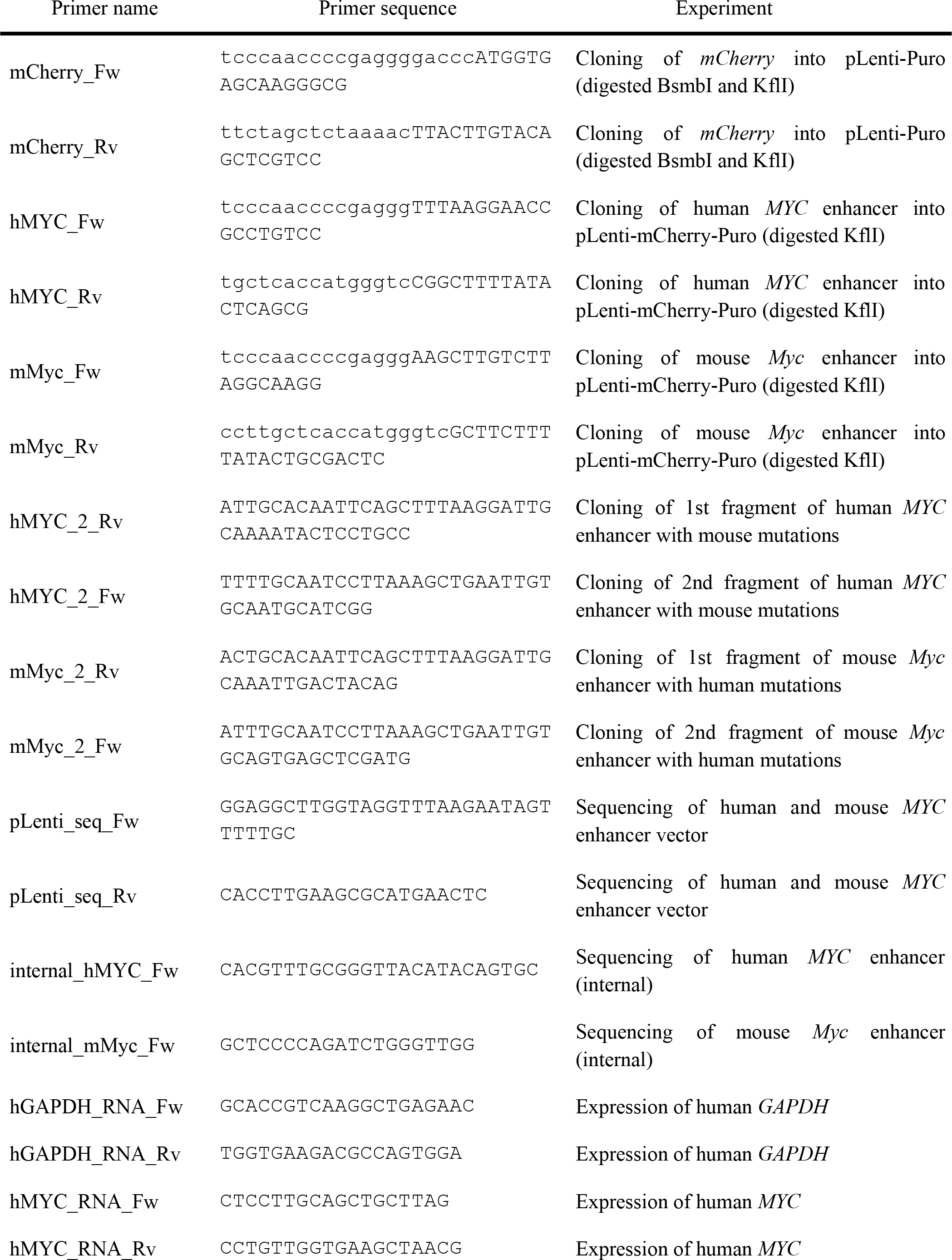
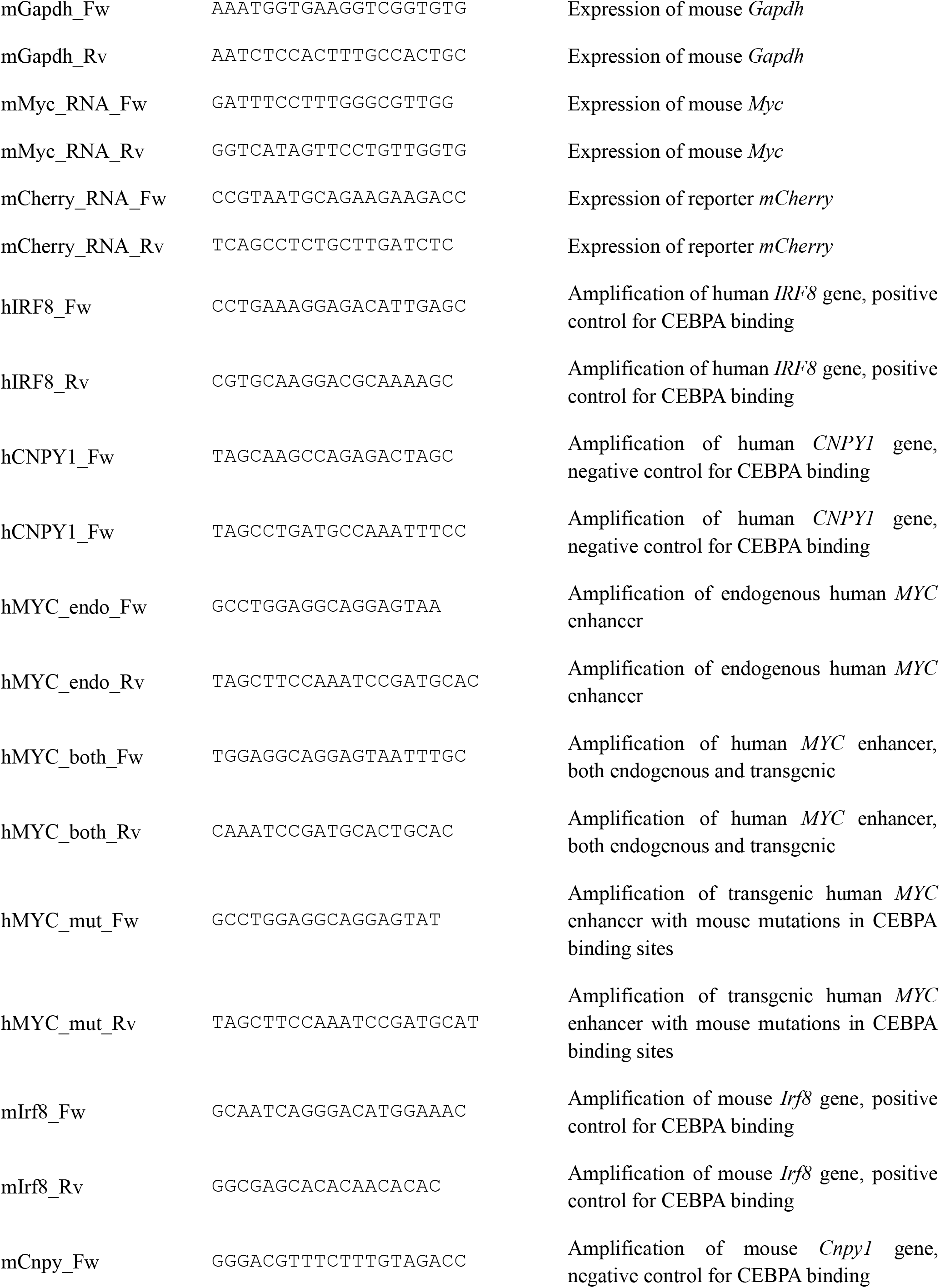
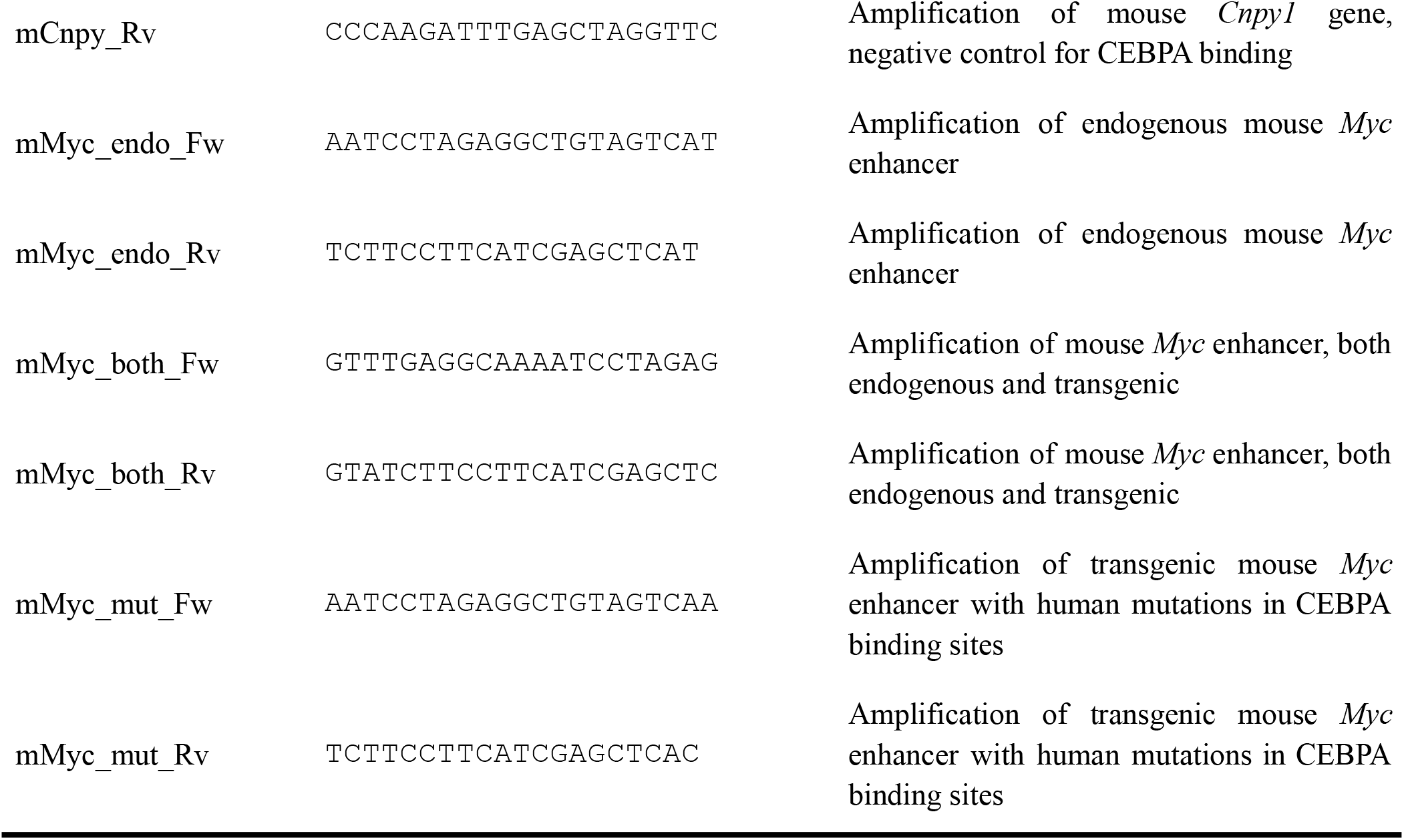
Primers used for qPCR, cloning and Sanger sequencing.

